# Robust eIF4B levels undermine invasive growth and immune evasion mechanisms in murine triple negative breast cancer models

**DOI:** 10.1101/2022.09.16.508318

**Authors:** Randall Smith, Leila Zabihi Diba, Aravind Srinivasan, Robert Zollo, Thomas Ossevoort, Qian He, Sean H. Colligan, Melissa Dolan, Yeshwanth Vedire, Tomina Sultana, Meera Venkatesh, Aayush P. Arora, Sarah Gawlak, Deschana Washington, Craig M. Brackett, Song Yao, John M.L. Ebos, Scott I. Abrams, Joseph Barbi, Sarah E. Walker

**Author notes:** Sarah E. Walker and Joseph Barbi (corresponding authors) and. These authors contributed equally (Leila Zabihi Diba, Aravind Srinivasan, Robert Zollo). **Author Contributions:** SEW conceived the project; SEW and JB designed, supervised, and financially supported all experiments described, analyzed results, and prepared the manuscript with input from all other authors. RJS assisted in experimental design and oversaw mouse model work and in vitro experiments. LZD, SEW, and AS made foundational observations. RZ, DW, and RJS performed flow cytometry analysis on samples from murine tumor experiments. TO, QH, AS, SG, TS, MV, APA, and LZD performed immunoblot analysis, cell culture, and in vitro studies. SC, MD, and YV assisted in the design and execution of mouse tumor studies. SY assisted in survival and gene expression analysis. CB, SA, and JLME provided key cell lines, reagents, and guidance pertaining to murine breast cancer models. All authors reviewed the manuscript and assisted in its preparation. **Competing Interest Statement:** The authors declare that no competing interests exist.

## Abstract

Dysregulated protein synthesis is seen in many aggressive cancers, including metastatic breast cancer. However, the specific contributions of certain translation initiation factors to in vivo disease remain undefined. This is particularly true of eIF4B, an RNA-binding protein and cofactor of the RNA helicase eIF4A and associated eIF4F cap-binding complex. While eIF4A, eIF4G, and eIF4E are well-known to contribute to the progression of many cancer types including metastatic breast cancers, the role played by eIF4B in breast cancer remains relatively unclear. We therefore explored how naturally divergent and experimentally modulated eIF4B levels impact tumor growth and progression in well-characterized murine triple negative breast cancer (TNBC) models. Surprisingly, we found that higher eIF4B levels in mouse and human breast cancers were associated with less aggressive phenotypes. shRNA-mediated eIF4B knockdown in TNBC lines failed to markedly alter proliferation and global translation in the cells in vitro and only modestly hindered their growth as primary mammary tumors growth in mice. However, eIF4B knockdown significantly enhanced invasive growth in vitro and exacerbated both tumor burden and mortality relative to nontargeting shRNA controls in a model of metastatic disease. Analysis of eIF4B levels and breast cancer patient survival reinforced a link to better outcomes. Interestingly, low eIF4B expression was also associated with more formidable immune evasion in vitro and in vivo, implicating a novel immunomodulatory role for this factor in the malignant setting that suggests a mode of action beyond its historical role as a co-activator of eIF4A/F.

**Significance Statement:** Metastasis is the leading cause of cancer-related mortality. Despite many advances in our understanding of this complex process and the molecular and cellular events involved, mechanisms that allow secondary tumors to arise and persist remain incompletely understood. Uncharacterized metastatic determinants active at the level of translational control may be exploitable as novel therapy targets or biomarkers predicting a tumor’s potential for spread and recurrence. Here we describe previously unrecognized consequences of dysregulated eIF4B levels in murine breast cancer that shed light on how this translation initiation factor contributes to disease outcomes. Our findings suggest that eIF4B levels direct metastatic risk and immune evasion, and further study should establish its value in personalized treatment decisions and development of future therapies.

## Introduction

Metastatic disease is the leading cause of patient mortality across most diverse cancer types. Exemplifying this is metastatic breast cancer, a deadly malignancy that presents a significant treatment challenge. While the 5-year survival rates for patients with *in situ* breast cancer is over 90%, the prospects for patients with metastatic disease are far more bleak (1, 2). There is mounting evidence that dysregulated protein synthesis plays a role in the onset and progression of aggressive cancers, including metastatic breast cancers (3). Several proteomic studies comparing breast cancer lines and samples with high and low metastatic potential (4–6) and studies of primary and metastatic tumors (7) alike have highlighted extensive protein-level distinctions associated with metastatic behavior in vivo. Furthermore, a number of the factors governing initiation of protein synthesis, a rate-limiting step that is critical for determining which proteins are made, are known to be dysregulated in breast and other cancers (8–10). This suggests that uncharacterized metastatic drivers and regulators may be operative at the level of translational control.

Translation begins with the formation of a 43S preinitiation complex (PIC), comprised of the 40S ribosomal subunit bound to several eukaryotic translation initiation factors and the initiator tRNA. Loading of this PIC and subsequent directional movement to locate the start codon, termed scanning, require the disruption of mRNA structures in the 5’ untranslated regions (UTRs) by the eIF4F complex, eIF4B, and DDX3 in mammals (11–13). When the PIC reaches a start codon in a good sequence context, rearrangements of the complex and its constituents occur, which arrest the 40S subunit in a scanning incompetent state with a fully engaged tRNA and allow large ribosomal subunit joining and subsequent peptide chain elongation (11, 14–16).

Translation efficiency and the extent to which individual transcripts rely on each eIF for translation depends on the length and degree of secondary structure in the 5’UTR and open reading frame (9, 17–19). Transcriptome-wide analyses of polysome-associated mRNAs have indicated that transcripts with hyper-dependence on eIF4A and eIF4B (yeast and mammalian proteins) have longer 5’UTRs and greater structural content (17, 18, 20). This likely stems from the ability of eIF4A to unwind secondary structures that obscure ribosome loading and scanning along the mRNA to the start codon(11). The RNA helicase activity of eIF4A is stimulated by eIF4B, eIF4G, and eIF4H; which promote a closed conformation of eIF4A that can bind to RNA; and it is inhibited by PDCD4 (11, 21, 22).

The critical importance of these events in the cancer setting is becoming increasingly clear. Environmental stresses present in the tumor microenvironment (i.e., hypoxia, nutrient deprivation) are capable of disrupting eIF4 factor activity and resulting translation output by altered posttranslational modifications and other mechanisms (23–25). eIF4B activity in particular is modulated by numerous cancer-relevant kinases, including p70/S6K (26–28). Recently, instances of collective and individual up-regulation of eIF4A, eIF4B, and eIF4E have been reported in cancers (29–31) with over-expression being associated with poor prognosis and metastatic disease (3). One study in particular noted this linkage for patients with endocrine receptor-negative breast cancer (9). Notably, its authors showed that experimental silencing of eIF4A in a human breast cancer cell line led to dramatic changes in gene expression and a less invasive in vitro growth phenotype (9). Similarly, eIF4E overexpression has been linked to pro-tumor processes like angiogenesis, metastasis (32), and worse patient outcomes in breast cancer (33). These findings strongly suggest alterations in eIF4-dependent translational control can have significant effects on the progression and spread of breast cancers, and ultimately patient outcomes. In contrast, the consequences of dysregulating eIF4B in mammalian cells remains less understood.

eIF4B is widely regarded as an accessory factor in the molecular decision making that governs initiation of protein synthesis. Capable of interactions with both mRNA and elements of the ribosome (34, 35), it plays an important but surprisingly ill-defined coordinating role in the process. eIF4B is best known for promoting the helicase activity of eIF4A to stimulate translation of mRNA transcripts with highly structured elements in their 5’UTRs (11, 13, 36). Since examples of eIF4A-reliant genes include several well-known, pro-tumor transcripts encoding proteins capable of fueling cancer cell division and aggressive tumor growth phenotypes (e.g., Cyclin D1, Cyclin D2 and CDK6 (17), Myc, notch1, MDM2, and Bcl2 (37)), it has been thought that up-regulation of eIF4B in breast cancer cells contributes to disease severity by promoting this decidedly pro-tumor translation program directed by eIF4A (9, 38, 39).

Yet, mounting lines of evidence suggest a more complex role for eIF4B in both translational control and human cancers. For example, insights recently gleaned from studies of eukaryotic translation in yeast show that eIF4B promotes translation transcripts with anticorrelated reliance on eIF4G, even though both factors stimulate eIF4A activity (18). This suggests that either eIF4B stimulates translation of some mRNAs by additional, uncharacterized mechanisms, or that there are multiple mechanisms by which eIF4A activity can be activated for distinct eIF4B and eIF4G-hyperdependent transcripts. Also, over-expression of eIF4B in mammalian cells can have opposing effects on translation depending on cell type and the levels of over-expression (40). Despite these observations, and the noted up-regulation of eIF4B levels in patients with breast (9, 38) and other cancers (39, 41), the precise in vivo consequences of this factor’s dysregulation in breast are not understood.

To shed light on eIF4B’s role in breast cancer, we scrutinized eIF4B expression patterns, patient survival data, and both the in vitro and in vivo results of shRNA-mediated knockdown od eIF4B in murine breast cancer models. Unexpectedly, we found an inverse association between eIF4B levels and the potential for aggressive disease progression and metastatic growth. While eIF4B knockdown modestly slowed the growth of primary tumors in vivo, invasive in vitro growth and the ability to form and grow as metastatic tumors were significantly increased with reduced eIF4B levels. Enhanced metastatic growth was linked in part to previously unappreciated relationships between eIF4B levels and mechanisms of immune evasion by tumors. Our results suggest that eIF4B plays an unexpectedly complex, stage-specific role in determining breast cancer outcomes. They may inform efforts to predict which tumors are likely to develop and progress as metastatic disease, and they beg further investigation of high- and low-eIF4B associated translatomes in breast cancers that could uncover novel regulators of metastatic growth and immune evasion.

## Results

### Low eIF4B levels are characteristic of aggressive breast cancers

Elements of protein synthesis control are known to be dysregulated in metastatic breast cancer (9) and other malignancies. The over-expression of some translation initiation factors (e.g., eIF4A/F, eIF4E) is well documented as is their ability to drive tumorigenesis, promote more aggressive phenotypes, and thereby contribute to poor patient survival (30, 38, 39, 42). In contrast, the degree and consequences of dysregulated eIF4B levels in such malignancies have received much less attention to date. Publicly available gene expression data (43, 44) reveal that a range of eIF4B transcript levels can be found among breast cancers, and the expression of this factor is generally lower in breast cancer tissues and cell lines than normal breast tissue and non-malignant cells (**Fig. 1A,B**) (45). Interestingly, when breast cancer subtypes were compared, eIF4B levels appeared *lowest* in triple negative breast cancers (TNBCs; **Fig. 1C**), known for their aggressive growth, resistance to treatments, and their propensity to spread and recur (46). Strikingly, eIF4B levels also decreased with increasing tumor grade (**Fig. 1D**) and a similar (albeit less pronounced) relationship was seen between eIF4B mRNA levels and disease stage data from the Cancer Genome Atlas (TCGA; **Fig. S2**). These observations linked robust expression of this initiation factor with early-stage, differentiated tumors with a lower capacity for metastatic spread. Such trends were distinct from the patterns of eIF4A expression in the same datasets, which were similar between cancerous and normal samples, highest in TNBC and high-grade tumors (3) showing no clear association with stage. Meanwhile eIF4G and eIF4E were upregulated in breast cancer tissues and lines, and their levels were positively related to tumor grade without clear relationship to subtype or stage (**Fig. S1A-C**). Thus, in striking contrast to other eIF4 factors whose levels have been widely associated with tumor-promoting mechanisms and disease progression, *low* eIF4B levels appears uniquely linked to negative disease attributes in breast cancer.

**Figure 1.**
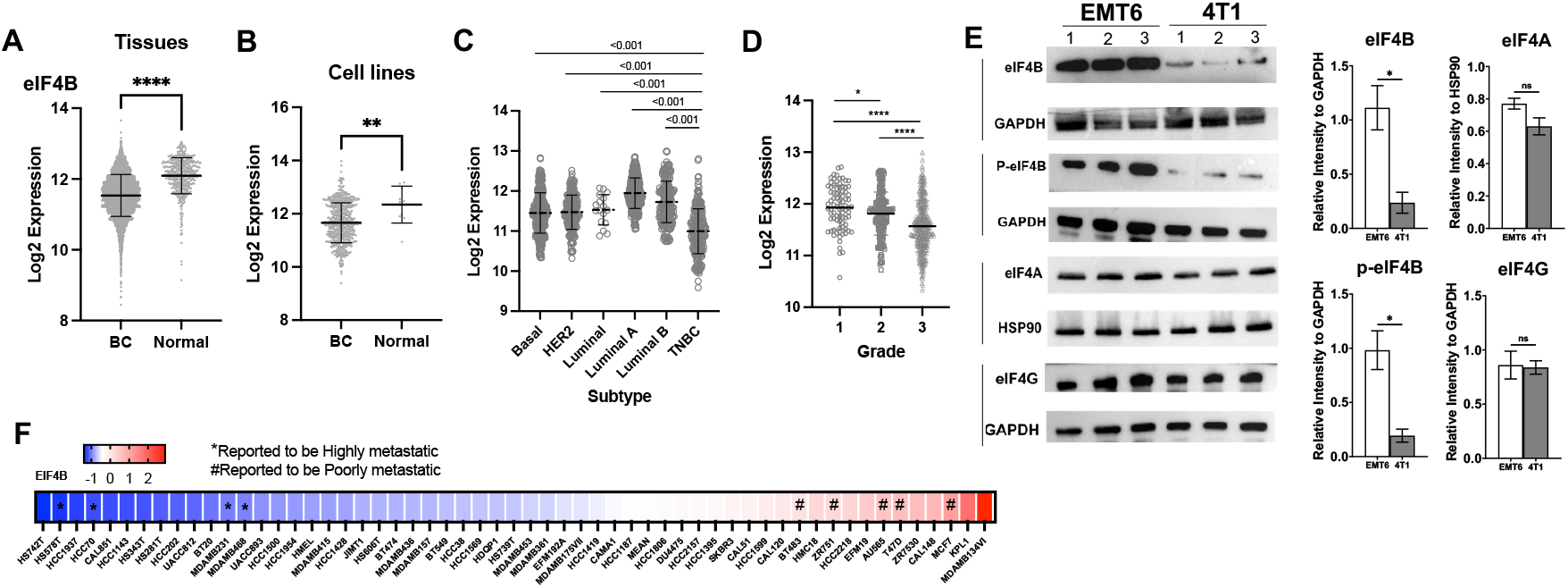
eIF4B expression levels vary according to breast cancer subtype, grade, and stage and are inversely related to reported metastatic tendencies. Gene expression (GeneChip microarray) datasets were accessed through the GENT2 analysis webtool. Levels of eIF4B transcript were compared in **(A)** breast cancer (BC) and normal breast tissue samples (n = 5574, and 475 respectively) as well as BC cell lines and non-malignant cell lines (n = 643, and 10 respectively). Similarly, eIF4B levels in breast cancers of distinct subtype **(C)** and tumor grades **(D)** were plotted and compared. **(E)** Protein levels of the indicated eIF4 factor in the murine TNBC lines EMT6 and 4T1 were determined by Western blotting. Cell lysates of each parental line (lanes1) and distinct subcultures initiated from in vivo isolates (lanes 2 and 3) were probed along with HSP90 and GAPDH (loading controls). Relative band densities normalized to loading controls are shown at right. **(F)** Relative eIF4B mRNA levels across human breast cancer line entries in the Cancer Cell Line Encyclopedia (CCLE) were found using CBioPortal and represented by a heat map. Cell lines reported to be highly (*) or poorly metastatic (#) are indicated. Panel E is representative of at least three independent experiments. Panels A-D depict the mean values (bars) for each group +/-SEM. Statistical differences (by t test) are shown (*p < 0.05, **p<0.01, ***p<0.001).

### TNBC lines with discrete metastatic potential exhibit distinct levels of translation initiation factors

To further investigate the relationship between eIF4B and the aggressiveness and metastatic potential of breast cancer, we surveyed the expression of several translation initiation factors in well-characterized murine TNBC cell lines. For this we chose two lines of common genetic background widely-used to model disease at opposite ends of the invasiveness spectrum: namely the highly metastatic 4T1 line and EMT6 cells, which are much less invasive with a generally poor capacity to form successful secondary tumors (47, 48). Strikingly, we found that EMT6 cells had markedly higher levels of eIF4B protein detectable by immunoblot analysis compared to their 4T1 counterparts, a trend also seen for activated eIF4B species phosphorylated at Serine 422 (27) (**Fig. 1E**). Rather surprisingly, no discernable differences were seen in these cells regarding eIF4A or eIF4G (**Fig. 1E**), factors generally regarded as eIF4B’s functional partners in promoting cap-dependent translation (11, 36). In addition to comparisons of parental lines (lanes 1), corroborating results were also seen in derivative subcultures generated from tumors recovered from the lung and livers of implanted syngeneic mice (essentially biological replicates, lanes 2 and 3). Thus, an over-abundance of eIF4B protein was linked to a *less aggressive* and *poorly metastatic phenotype* in murine breast cancer models. This relationship was also evident in *human* breast cancer cell line gene expression data found in the Cell line Encyclopedia (CCLE) (49). Entries with well-known metastatic tendencies including the highly invasive MDA-MB-231 (50, 51) and Hs578T (52) lines were among the lowest relative eIF4B expressors. In contrast, MCF7, AU565, ZR751, BT-483, and T47D, which are generally described as poorly metastatic in vivo (53–55), were relatively high in their eIF4B levels (**Fig. 1F**). This apparent inverse relationship between eIF4B and metastatic capacity in murine and human breast cancers supports a previously unexplored role of this translation initiation factor in limiting invasive growth in breast cancers when expressed in abundance.

### eIF4B knockdown in TNBC cells promotes invasive growth in vitro

The implications of differential eIF4B expression in breast cancer remain largely unknown. To address this gap in knowledge, we used lentiviral-delivered eIF4B-silencing (sh-eIF4B) and non-targeting control (sh-sc) constructs to generate stable knockdown and control lines from parental EMT6 and 4T1 cells. Effective, but incomplete eIF4B knockdown capacity was confirmed for three separate constructs (**Fig. S3A**), and these lines were subjected to several in vitro assays. In contrast to a prior study that reported dramatic proliferation and survival defects upon inducible (and near complete) eIF4B knockdown in a human breast cancer line (56), we observed no marked deficiency in growth rate, cell cycle progression, or viability upon sh-eIF4B under normal 2-dimensional cell culture conditions (**Fig. S3C-E**). In fact, a slight growth advantage for some knockdown lines was noted (**Fig. S3C**). Rather unexpectedly, given its well-known participation in the molecular events at the start of protein synthesis, sh-eIF4B and control EMT6 lines behaved similarly in an assay of translation output relying on the detection of surface proteins containing incorporated puromycin (i.e., a SUnSET assay)(57), and only a modest defect was seen with knockdown in the 4T1 line (**Fig. S3F**).

Similarly, a widely-used wound-healing assay of cell motility and growth in 2-D culture revealed no significant differences in the rate or ultimate success of wound closure by sh-sc and sh-4b derivatives of either EMT6 or 4T1 line over a 24 (**Fig. S4A, B**) or 48-hour observation period (data not shown). This aligns with a recent study showing several invasive ductal breast cancer cell lines (characterized elsewhere as having high or low relative eIF4B levels) can readily close the gap in this assay (53). Together these results suggest that partial eIF4B silencing does not generally impair translation or core cellular processes in breast cancer cells in vitro.

To gain insight into the relationship between eIF4B levels and the capacity for invasive growth, we compared the ability of the aforementioned cell lines to invade and transmigrate a simulated extracellular matrix (ECM). Here, the ability of fluorescently-labeled cells to traverse a porous, ECM-coated transwell insert and emerge on the undersurface of the resulting barrier in response to a serum gradient was observed over time with automated digital fluorescence microscopy. In keeping with their aggressive reputation, seeded 4T1 cells rapidly migrated through the ECM barrier, with a non-trivial degree of migration seen even in the absence of serum attractant. However, the knockdown of eIF4B in either the EMT6 or 4T1 line dramatically enhanced the degree of invasive migration over each scrambled control (**Fig. 2 A,B**). Microscopic examination of sh-eIF4B harboring cells cultured on ECM also revealed indications of increased focal adhesion points with the simulated tissue. This latter in vitro behavior is strongly associated with enhanced integrin engagement and a high potential for cancer cell invasiveness and metastatic growth in vivo (58, 59). Specifically, increased cell perimeter and reciprocally reduced circularity was seen upon eIF4B knockdown relative to controls for both lines, while divergent effects on cell surface area were noted in EMT6 and 4T1 lines (**Fig. 2C, D**). These findings indicate a previously unreported role for eIF4B in moderating invasion that may have implications for the growth and dissemination of tumors in vivo.

**Figure 2.**
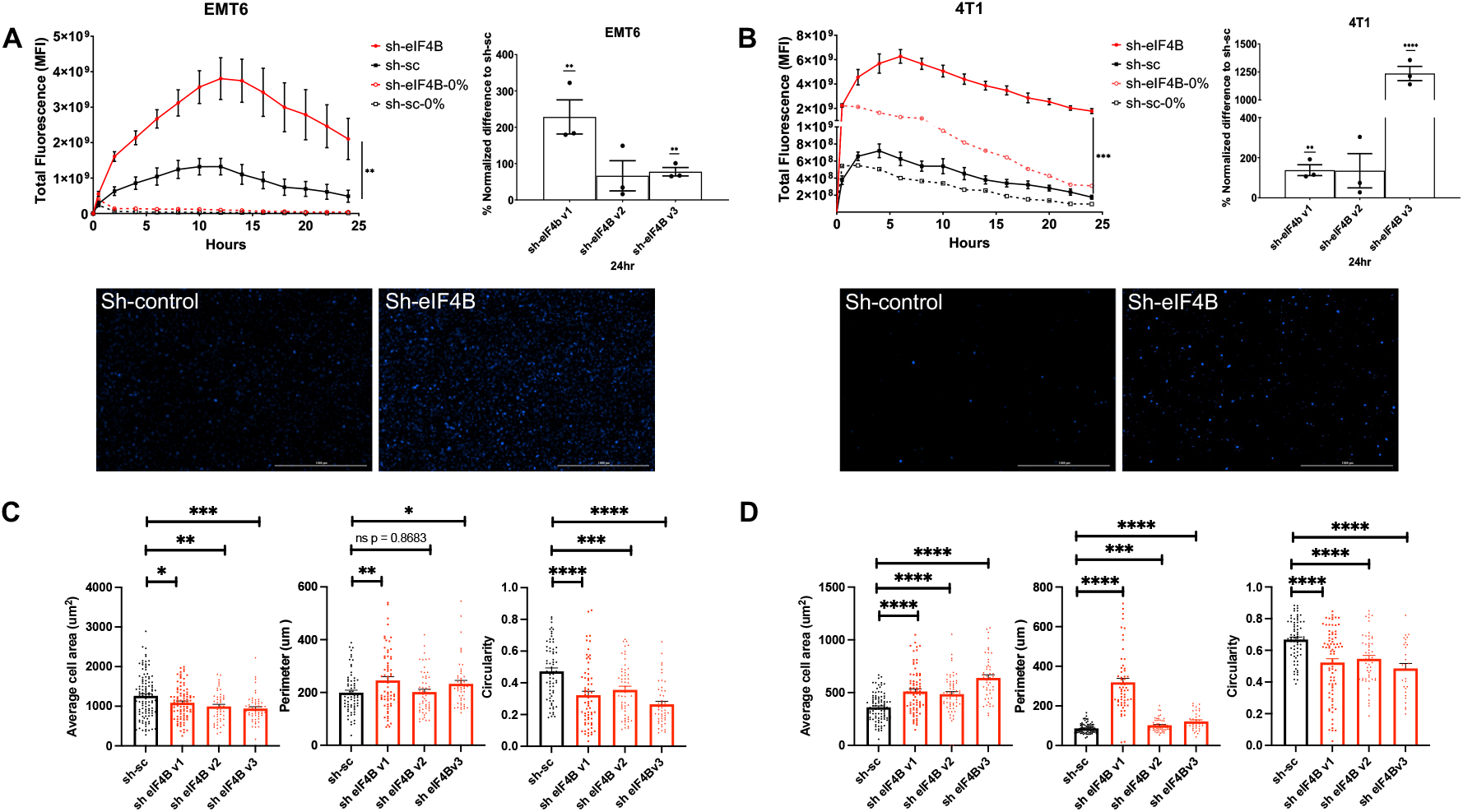
Invasive growth by murine breast cancer lines in vitro is enhanced by eIF4B knockdown. **(A,B)** EMT6/4T1 shRNA knockdown (red lines) and control lines (black lines) were pre-stained with cell trace violet. Cells were added on top of pre-coated (Geltrex-ECM) transwell membrane inserts and RPMI media with 10% FBS (solid lines) or 0% FBS (dashed lines) was added below the inserts. Real-time imaging was performed over 24hrs and mean fluorescent intensity of the cell trace violet recorded at 2-hour intervals. Line graphs depic kinetics over 24hrs, with peak migration occurring at 6 (4T1) and 12 (EMT6) hours. Bar graphs depict MFI at 24hrs. Shown are representative fluorescence microscopy images and the mean fluorescence intensities (MFIs; +/-SEM) from one of at least three independent experiments. **(C,D)** The indicated cell lines were grown on simulated ECM (Geltrex coated 24 well plates) for 12 hours, fixed, and stained with Alexa Fluor 488 conjugated actin/phalloidin and nuclei counterstained with Hoescht dye, imaged, and their morphology (area, circularity, and perimeter) was assessed and quantified using image J software built in tools. Cells (n>100) from 10 fields of view over 3 wells and 3 independent experimental replicates were used. Statistical differences (by t test) are shown (*p < 0.05, **p<0.01, ***p<0.001, ****p< 0.0001).

### eIF4B knockdown enhances the in vivo metastatic potential of murine breast cancer lines

eIF4B upregulation in breast tumors has been reported with apparent pro-tumor effects in vitro (9, 38). Yet the specific contributions of this factor’s dysregulation to in vivo tumor growth and the potential for metastatic progression have gone largely unexplored. This, coupled with the robust expression of eIF4B by less aggressive murine and human breast cancer lines and enhanced invasion seen upon eIF4B knockdown in vitro led us to explore the impact of differential eIF4B level on mammary tumor growth and disease in vivo. To this end, eIF4B knockdown and control lines of 4T1 and EMT6 cells were generated that also harbored a luciferase reporter (4T1-luc, EMT6-luc) (**Fig. S3A**). These lines were implanted into the mammary fat pads of female BALB/c mice, and the growth of primary tumors was monitored over time by digital caliper measurements and at endpoint using bioluminescence imagery (BLI). Not surprisingly, implanted 4T1 control cells rapidly developed into large primary tumors while EMT6 tumors progressed relatively slowly. In both models, eIF4B knockdown resulted in a modest delay in mammary tumor growth relative to controls (**Fig. 3A**), suggesting the initiation factor contributes marginally to the development of primary breast tumors.

**Figure 3.**
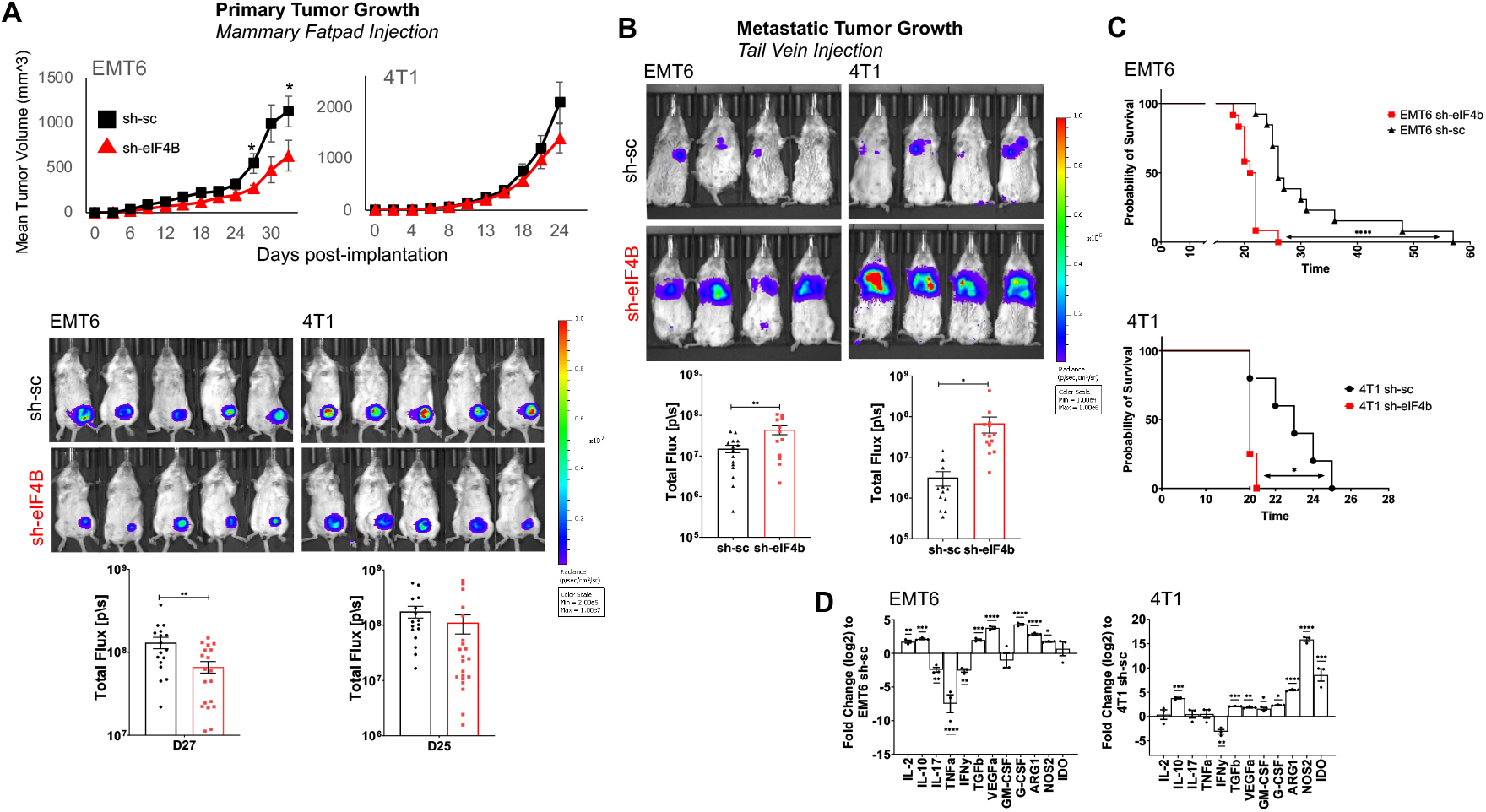
Downmodulation of eIF4B levels slows primary tumor growth but markedly accelerates metastatic disease in mice. Knockdown of 4B in luciferase-expressing 4T1 and EMT6 breast cancer cell lines was achieved by lentiviral delivery of sh-RNA (sh-eIF4B, 4B) and non-targeting control (ctrl, sh-c) constructs. The effect of eIF4B knockdown on primary tumor growth was assessed after implantation of Balb/c mice with 5×10^4^ cells by intra-mammary fat pad (i.m.) injection (n=7-8/group). **(A)** i.m. tumor development was monitored by caliper measurements (upper panel) and in vivo bioluminescence (IVIS) after injection of d-luciferin (lower panel). **(B)** Effects of eIF4B knockdown on the formation and progression of lung metastases was determined by intravenous (i.v.) injection of Balb/c mice (n=5-7/group) with 5×10^4^ cells cells. Here tumor burden was assessed by IVIS 26 days. Representative IVIS images (left) and mean tumor burdens +/-SEM (right) are shown. **(C)** The survival of recipient BALB/c mice challenged in panel B were compared in Kaplan Meier plots. **(D)** Gene expression analysis (qRT-PCR) of tumor bearing lungs from indicated lines. Expression was quantified as the fold change (log2) to sh-sc controls per cell line. Statistical differences (by t test) are shown (*p < 0.05, **p<0.01, ***p<0.001, ****p< 0.0001).

To understand the impact of experimentally diminished eIF4B levels on the establishment of metastatic tumors independent of primary tumor growth effects, we utilized an i.v. challenge approach used to model hematological spread and growth of pulmonary metastases. Here we injected eIF4B knockdown and control EMT6-luc lines via the tail vein, and monitored both the development of tumor burden (luciferase signal), and the survival of recipient mice post-challenge. While poorly capable of forming spontaneous lung metastases in immunocompetent mice, EMT6-luc cells readily give rise to pulmonary tumors when injected i.v. into naïve BALB/c mice (48). In line with this observation, both lines also gave rise to nascent lung metastases evident in a colony forming assay within four days post-injection (**Fig. S5A**) and established pulmonary tumor burden within 26 days, detectable by BLI (**Fig. 3B**). Notably, recipients of sh-eIF4B-luc cells developed significantly more prominent lung tumor burdens than mice challenged with sh-control cells of either line (**Fig. 3B**. Strikingly, the eIF4B knockdown groups also displayed significantly shorter survival times compared to controls (**Fig. 3C**). This result suggests that eIF4B may play an unexpected role limiting the development and progression of metastatic tumors through yet undefined mechanisms.

Repeating these experiments in syngeneic, immunodeficient mice allowed us to assess the involvement of the immune defenses in these effects of eIF4B knockdown. Here, Rag2-deficient BALB/c mice also lacking expression of the IL-2receptor gamma subunit (Rag2-/-,γc-/-mice, which lack functional T, B, and NK immune cell populations) received sh-eIF4B and sh-control EMT6-luc and 4T1-luc cells i.v. Not surprisingly, these mice developed established lung tumors as early as seven days post injection (**Fig. S5B, C**), and mortality was seen with generally accelerated kinetics in both models (**Fig. S5D**). As in immune-competent mice, sh-eIF4B cells gave rise to higher lung tumor burdens than control lines, though the effect in this context was modest (**Fig. S5B, C**). Notably, the enhanced mortality resulting from eIF4B knockdown that was clear in immunocompetent animals was not seen in the Rag2-/-,γc-/-mice (**Fig. S5D**) suggesting a potential role for the immune system in determining at least some of the in vivo consequences of eIF4B dysregulation. In line with this notion, evidence of decidedly pro-tumor immune modulation was seen in the lung metastatic niche. Specifically, transcripts encoding the tumoricidal cytokine IFNγ and other proinflammatory cytokines were downregulated upon eIF4B knockdown, while mediators of immune suppression (e.g., IL-10, TGF-beta, and iNOS) were apparently enhanced (**Fig. 3D**). Together these results suggest that the enhanced metastatic growth seen when eIF4B levels are reduced stem from immune evasion of those cells.

### eIF4B knockdown in murine TNBC is associated with immune dysfunction in vitro and in vivo

The persistence and growth of disseminated cancer cells and nascent secondary tumors depend in part on the ability to evade detection and elimination by the immune system (60). Indeed, several mechanisms of immune suppression have been shown to promote the growth of secondary breast tumors including the induction and mobilization of suppressor cell types including Regulatory T cells (Tregs) and myeloid derived suppressor cells (MDSCs), and the engagement of inhibitory signaling pathways such as the PD1:PD-L1 axis (60, 61). It was recently shown that even breast cancer lines with low invasive growth capacity (i.e., EMT6) can disseminate and seed secondary tumors in vivo. Yet successful metastatic growth is undercut by the immune defenses, CD8+ T cells in particular. Meanwhile, enhancement of suppressor cell pools can permit metastatic growth even in this less aggressive model (48).

To determine whether tumor eIF4B levels influence metastatic growth by modulating anti-tumor immunity, we characterized the leukocyte presence and disposition in the tumor microenvironment (TME) generated in our in vivo studies. Multicolor flow cytometry analysis of tumor bearing lungs 26 days post-i.v. injection revealed a dearth of T cells in the TME upon eIF4B knockdown. This was particularly true for non-regulatory or conventional T cells, “Tcon”, which were less frequent among the tumor associated leukocytes **(Fig. S6A left, S7A**). These cells also appeared less proliferative (indicated by reduced levels of KI67), especially in the 4T1 model (**Fig. S6A right**). Moreover, these cells were less committed to an activated effector phenotype in the sh-eIF4B groups of both models as evidenced by the reduced frequencies of CD44^high^/CD62L^low^ and the mean fluorescence intensity (MFI) of the activation marker CD44 (**Fig. S6B, C**). Activated effector CD8+ (eCD8+) T cell pools were also compromised in lungs harboring sh-eIF4B tumor. These potential cytotoxic tumor cells comprised a smaller fraction of both the CD8+ and overall leukocyte populations (**Fig. S6D**), and their expression of the activation marker CD44 was reduced upon eIF4B knockdown (**Fig. S6E**). Importantly, the relative abundance of suppressive Regulatory T cells (Tregs) with an activated phenotype (eTregs) to these effector CD8+ T cells (i.e., the eTreg:eCD8 ratio) was elevated when tumor cells harbored reduced eIF4B levels (**Fig. S6F**). Thus, reduced eIF4B expression was linked to indications of dampened anti-tumor immunity.

The effects of eIF4B knockdown on the bulk frequency of tumor-associated Foxp3+ Tregs in this model of metastatic disease were distinct in the cell lines examined (increased in EMT6, decreased in 4T1) (**Fig. S7A**). However, these suppressor cells consistently displayed higher levels of CD44 and immune checkpoint factors CTLA-4, PD-1, and PD-L1 in mice injected with sh-eIF4B lines (**Fig. S7B-E**). Since these markers are typically upregulated by activated Tregs and those accumulating within the TME (62–64), these findings associate lower eIF4B levels with the activity of a major tumor-relevant suppressor cell type and potential immune dysfunction in the metastatic tumor niche.

Interestingly, eIF4B knockdown in primary mammary fat pad tumors resulted in heightened proportions of tumor infiltrating leukocytes (TILs, CD45+) **(Fig. S8A**). Yet, among these cells, Tcon frequencies were reduced compared to controls with evidence of dysfunction (i.e., upregulation of the immune checkpoint factors TIM3 and PD-1; **Fig. S8B-D**). The CD8+ T-cell compartment was also compromised among the TILs of implanted sh-eIF4B tumors, showing signs of lower viability and robust expression of Lag3, PD-1, and TIM3 (**Fig. S8E-H**). Treg proportions were higher among the CD4+ TILs of the sh-eIF4B groups, and these cells were increasingly positive for Lag3 (the 4T1 line in particular) and PD-L1 compared to controls (**Fig. S9A-C**). These findings suggest that some of the immunological effects resulting in dampened intra-tumoral immune activity are operative across disease models.

A recent study suggested that the differential invasiveness seen between EMT6 and 4T1 lines at least partly reflects their divergent capacities to induce MDSCs (47). Indeed, cells of this lineage are known to aid tumor growth and metastasis by opposing the activity of anti-tumor effector cells, conditioning the pre-metastatic niche, and promoting angiogenesis (65). Expectedly, we found that 4T1 tumor-associated tissues contained greater frequencies of CD11b+/GR1+ cells (likely MDSCs) than those in the EMT6 model. In the former, eIF4B knockdown increased MDSC proportions within the tumor associated myeloid compartment without a discernable effect on these cells in the latter (**Fig. S10 A, B**). These results suggest that when prominently induced by line-specific factors, MDSCs may be enhanced under low-eIF4B conditions.

Collectively, these findings suggest that lower eIF4B levels may support mechanisms for dampening immune activity needed for an effective antitumor response. To directly observe how TNBC cells with differential eIF4B expression modulate leukocytes activity, a tumor-splenocyte co-culture system was used (**Fig. S11A**). The expansion of dividing CD8 T cells of an activated/effector phenotype (identified by dilution of cell-trace violet dye and high surface levels of CD44) was inhibited by the presence of co-cultured TNBC cells after 48 hours of stimulation. This suppression of potential anti-tumor effectors was significantly more pronounced when eIF4B was knocked down in either line (**Fig. S11A**). Similarly, the overall frequency of effector CD8+ T cells were less elevated by in vitro activation in samples with reduced eIF4B levels (**Fig. S11B**). In contrast, PD-1 expression by co-cultured CD4+ and both the division and frequencies of activated Tregs were upregulated by stimulation in the presence of knockdown lines (**Fig. S11C-E**). These results demonstrate that lowering eIF4B levels can amplify tumor cell-associated inhibition of T cell activation while promoting mediators of immune suppression – effects that would be expected to improve immune evasion in vivo.

### eIF4B levels are inversely related to baseline and induced PD-L1 expression by TNBC cells

Disseminated tumor cells and micro-metastases can persist and thrive in the face of immune surveillance by upregulating immune checkpoint signaling pathways such as PD1/PD-L1. Importantly, the upregulation of PD-L1 by tumor cells was recently shown to be controlled at the translation level by the eIF4A/eIF4F-dependent expression of STAT1 (66). However, the involvement of eIF4B in this translationally governed process has not been explored.

When we examined the effects of shRNA-mediated eIF4B knockdown on the ability to induce PD-L1, as expected (66), IFNγ exposure greatly up-regulated PD-L1 signal in both lines in vitro. However, rather surprisingly given eIF4B’s widely accepted role as a co-activator of eIF4A, both the percentage of PD-L1-expressing cells and their intensity of PD-L1 expression were *enhanced* upon down-modulation eIF4B expression relative to controls (**Fig. 4A, Fig. S12 A**). This enhanced PD-L1 induction was accompanied by elevated STAT1 protein levels in both lines upon exposure to cytokine (**Fig. S12B**), a key mediator of IFNγ-driven PD-L1 upregulation by tumor cells (66). RT-PCR further confirmed the enhanced upregulation of PD-L1 and STAT1 coincident with eIF4B targeting (**Fig. S12C).** Strikingly, flow cytometric analysis of viable, non-immune (CD45-) cells of the tumor (predominantly tumor cells) in our i.m. and i.v. mouse model experiments also revealed substantial upregulation of PD-L1 upon eIF4B knockdown (**Fig. 4B, C**). In these experiments, a modest drop in tumor cell viability was seen when eIF4B expression was targeted in the presence of proinflammatory cytokine (**Fig. S12D**) and in vivo (**Fig. 4D**). This observation is in line with a previously described role for the factor in promoting survival in Hela (40) and breast cancer cell lines (56). Interestingly, low-eIF4B expressing TNBC cells in our in vitro co-culture assays also displayed heightened PD-L1 levels compared to controls, and these increased stepwise with co-cultured immune cells and T cell stimulating anti-CD3 antibodies (**Fig. 4E**). Since the PD-1:PD-L1 axis supports the conversion and maintenance of Tregs (67), the expanding pool of these cells that we see in our co-culture experiments could stem from elevated PD-L1 induction under eIF4B knockdown conditions.

**Figure 4.**
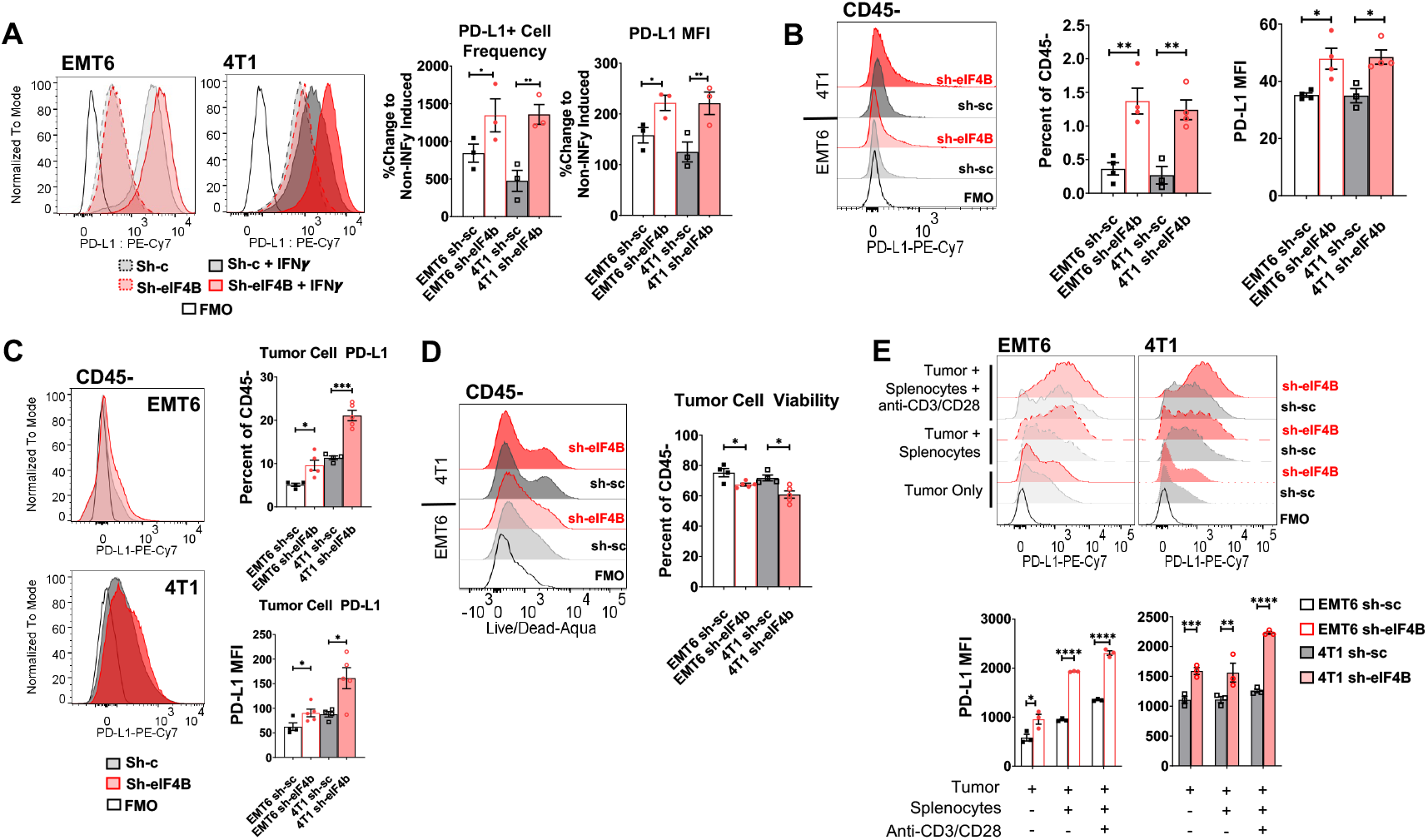
Lower levels of eIF4B favor baseline expression and IFNI driven upregulation of PD-L1 by murine breast cancer cells. Stable transductants of the EMT6 and 4T1 cell lines were generated with either control or sh-eIF4B constructs, grown in vitro and seeded in 6-well culture plates prior to exposed to recombinant IFNI (100ug/ml) or media alone for 24 hours. The frequency of cells expressing surface PD-L1 and the surface levels of PD-L1 on these cells (MFI) with and without cytokine treatment were determined by immunostaining followed by flow cytometry analysis **(A)**. Similarly, PD-L1 levels on non-immune (predominantly tumor) cells recovered from the tumor microenvironment in the i.v. **(B)** and i.m. **(C)** challenge models were found as well as the viability of tumor cells in vivo **(D)**. Levels of PD-L1 on tumor cells recovered from the indicated conditions in vitro coculture studies were also determined by flow cytometry **(E)**. Histogram overlays depict the results of 2-5 independent experiments, and the mean results (+/-SEM) of a representative trial are shown. (*p < 0.05, **p<0.01, ***p<0.001, ****p< 0.0001) by t test.

Analysis of eIF4B and PD-L1 expression (RNASeq) data available through TCGA further supported a significantly negative relationship between these factors. As shown in Fig. S13A, a clearly inverted association was seen between PD-L1 transcript and that of eIF4B that stands in contrast to eIF4A, eIF4G, eIF4E and eIF4H mRNA levels that showed either no clear relationship with PD-L1 expression, or a positive one (**Fig. S13B-E**). Interestingly, high (above median) expression of PDCD4, a known antagonist of eIF4A, was predictably linked to lower PD-L1 levels – an inverse relationship that was less pronounced than that between eIF4B and PD-L1 (**Fig. S13F**). In all, these observations suggest that heightened expression of eIF4B in breast cancer cells adversely affect a major mechanism of immune evasion, and this unappreciated effect may contribute to differential metastatic potential among mammary cancers, as well as resistance to endogenous and therapy-induced anti-tumor immunity. They also beg further exploration of eIF4B’s role in the metastatic process and patient outcomes.

### Evidence for a stage-specific association between eIF4B levels and patient survival outcomes

While our findings in the murine system suggest an unappreciated role for eIF4B in regulating metastatic growth and immune evasion, their direct implications for human disease are admittedly difficult to assign. Relative eIF4B protein levels (determined by immunohistochemistry) were previously associated with worse survival in ER-negative breast cancer patients (9). Interestingly, another study using mass spectrometry to characterize the global proteosome of TNBC patients (68) found an opposite albeit not statistically significant trend. Reanalysis of this data revealed upper quartile Overall Survival (OS) times for eIF4B-high and -low patient cohorts of 100 and 47 months, respectively (**Fig. S14**). This observation suggests a more complex relationship may exist between eIF4B and disease outcome in breast cancer. Analysis of breast cancer entries in TCGA suggested no clear relationship between eIF4B expression over all patients (**Fig. S15A**). However, separate analysis of early (stage I, II) and late-stage (stage III,IV) breast cancer patients revealed that high (above median) eIF4B levels are associated with a marginally negative effect among the former group (**Fig. S15B**; p=0.087) and a statistically significant benefit for the latter (**Fig. S15C**; p=0.047) (49). Similarly, the HRs calculated for these patient groups also suggested a shift in how eIF4B levels impact patient survival by stage (stage I, HR = 3.32). These results agree with those obtained in our mouse model experiments that suggest robust eIF4B levels may impart a pro-tumor advantage in primary tumors and a disadvantage in the metastatic growth phase characteristic of advanced stage disease.

A separate, compiled data set consisting of patient survival and eIF4B gene expression (microarray) data from 14 publicly-available studies, agnostic of treatment status, revealed significantly better survival rates for high (above median eIF4B expressors) compared to relatively low (below median) expressors (p=6.9×10^−6^; **Fig. S15D**). Interestingly, datasets from studies of predominately early stage patients (stage I and II) revealed an inverted trend, with high eIF4B-expressers displaying poorer overall survival (OS), though this was not statistically significant (GSE69031, (69); GSE58812(70)), an observation in line with a survival advantage of high eIF4B specifically in late stage disease. Recurrence free survival (RFS) and distant metastasis free survival (DMFS) were similarly better for high expressors of eIF4B compared to low expressers (p= 3.2×10^−7^ and 4.3×10^−10^, respectively; **Fig. S15E, F**), and comparison of post-progression survival (PPS) times for an admittedly smaller number of patient samples (n= 229/group) linked above-median eIF4B levels with statistically significant longer survival (p= 0.0432; **Fig. S15G**). Restricting analysis to data from patients having not received systemic therapies revealed significantly better RFS and DMFS outcomes (p= 0.0013 and 0.0024, respectively) and nominally improved OS and PPS (p= 0.3607 and 0.1905, respectively) for high expressors of eIF4B compared to low expressers (**Fig. S16A-D**). Suggesting a uniquely negative role for eIF4B in the metastatic growth phase of breast cancers, DMFS was less for patients with above median eIF4A and G and not clearly effected by levels of eIF4E and H (**Fig. S16E-H**). Separate analysis of lymph node (LN) positive breast cancers (i.e., those showing some metastatic spread) and LN-negative cases revealed similar associations with eIF4B and OS (**Fig. S17A**). However, the differences between relatively high and low eIF4B expressing patients in terms of RFS and DMFS were noticeably muted in LN-negative cases (**Fig. S17B**) – further linking low eIF4B levels with worse outcomes in metastatic disease.

In all, these results suggest that dysregulation of eIF4B in breast cancer can have unexpected, yet clinically relevant effects that are both unique among those of eIF4 and other initiation factors and important for metastatic disease and cancer-related mortality. They also suggest a previously unappreciated disease stage-specific relationship between eIF4B levels, invasive growth, immune evasion mechanisms, and disease outcome.

## Discussion

Disrupted control of protein synthesis has been seen in numerous disease states including human metastatic breast cancers. In contrast to other eIF4 factors that have received more investigative attention, the role played by eIF4B in breast cancer is largely unclear. Being a well-appreciated co-activator of eIF4A helicase activity, eIF4B is often thought to facilitate the oncogenic processes dependent on eIF4A/F-dependent translation, which includes cancer cell proliferation, EMT, resistance to apoptosis, glucose metabolism (71, 72), and, more recently, immune evasion (66). In line with a pro-tumor role for eIF4B, a number of largely in vitro studies have reported detrimental effects of eIF4B silencing on malignant cell survival, proliferation (9), and the ability to resist genotoxic stress (73). Yet, the consequences of modulating eIF4B levels on tumor progression in vivo are much less clear.

In the present study, we set out to explore the impact of dysregulated eIF4B levels at the complex in vivo intersection of tumor intrinsic and extrinsic factors that determine breast cancer disease outcomes. Assessing eIF4B expression patterns in breast cancer patient data, cell lines, and widely-used murine models revealed an unexpected association between robust eIF4B levels and less-aggressive cancers with reduced capacity for metastatic growth. In line with this, knocking down eIF4B in murine breast cancer models significantly accelerated progression of fatal metastatic disease with only modestly inhibition of tumor growth at the primary site. This surprising enhancement in metastatic growth was linked in part to a more invasive growth phenotype (observed in vitro) and improved immune evasion evident in multiple in vitro and in vivo assays. Supporting the notion that eIF4B plays distinct roles in the primary and secondary disease sites are the survival data of breast cancer patients with disparate tumor eIF4B levels. Taken together and along with previously described, pro-tumor effects attributed to eIF4B in breast and other cancer cells, our results suggest that eIF4B plays an unexpectedly complex, stage-specific role in determining breast cancer outcomes.

When eIF4B levels are experimentally reduced, implanted tumors in the mammary fat pad grow somewhat slower, and early-stage patients (typified by the presence of primary tumors with limited dissemination) fare marginally better than high eIF4B expressors. These observations likely stem from the pro-tumor roles played by eIF4B in promoting cell division and survival characterized in previous studies of eIF4B in cancer cell lines. For instance, Wang et al., showed that shRNA mediated knockdown of eIF4B in human breast cancer lines led to a pronounced defect in the growth rate and survival of cells in vitro (56). Modelska et al., also reported a slower in vitro growth rate for the human breast cancer cell line MCF7 upon eIF4B knockdown (9).

While these prior findings are compatible with our observations in mice with primary tumors and early-stage breast cancer patients, there is little in the published literature to suggest the surprisingly opposite role of eIF4B suggested by findings in the metastatic tumor model and advanced-stage patients. Levels of phosphorylated eIF4E are known to be elevated in early-stage carcinomas compared to late-stage cases (74), suggesting potential disease stage-specific activity of translation factors. Interestingly, and germane to our study, Hellwig et al., found *EIF4B* to be among four genes significantly associated with metastasis-free survival in breast cancer patients three-years or more after primary treatment (surgery) (75). In line with our analysis of public survival datasets, this supports a role for eIF4B as a negative regulator of metastasis and breast cancer progression at the advanced stage.

Recently, a computational study by Wu and Wagner assessed the relationship between translation factors and patient survival across all cancers within TGCA (76) and found a strong association between high eIF4G levels and poor prognosis. Higher eIF4B levels, on the other hand, were rather uniquely associated with better patient survival (76). Interestingly, despite Kaplan Meier analysis highlighting a marked survival advantage among high-eIF4B expressing patients, the statistical significance of this benefit was left unclear owing to violations of the study’s hazard ratio (HR) assumptions – which could have reflected an inconsistent directionality of the HR across cancer types or disease stages, as we have found for breast cancer (**Fig. S13**). As such, at the very least, this study supports the notion that eIF4B’s relationship with the survival of patients with breast, and potentially other, cancers can be distinct from other eIF family members.

For some time now, it has been appreciated that the relative level of eIF4B expression (i.e., the degree of over-expression) can have distinct consequences for mammalian translation. Some studies have shown an enhancement of cap-dependent translation in cells over-expressing eIF4B (77) while others found stunted protein synthesis (28, 78). Indeed, in 2006, Sonenberg and colleagues speculated that excess eIF4B levels could inhibit translation through the sequestration of other translation initiation factors. Such would be expected to disrupt the formation of functional complexes while generating potentially inactive pools of eIF4B-interacting partners such as eIF4A, eIF3, and small ribosomal subunits (27). In this study, we did not observe considerable effects on bulk translation or considerable differences in cellular fitness indicative of compromised protein synthesis upon partial eIF4B knockdown or when eIF4B levels were naturally high (though EMT6 cells displayed somewhat lower puromycin incorporation than 4T1 cells).

It is possible that our findings instead reflect a liberation of translation machinery needed for the synthesis of metastasis-promoting proteins, when translation of mRNAs requiring eIF4B activity are repressed. Such would be compatible with the more effective PD-L1 upregulation, a process driven by eIF4A/eIF4F activity, (66) which we saw upon eIF4B knockdown (**Fig. 4**). Previous results with purified human proteins suggest that binding of eIF4B and the related protein eIF4H to eIF4A are mutually exclusive(79). The eIF4H protein encompasses a similar core architecture to eIF4B but lacks N- and C-terminal elements responsible for interactions with eIF3, PABP, and possibly the 40S subunit, and sites subject to regulatory phosphorylation. It is possible that in a reduced eIF4B context, eIF4H binds to eIF4A more effectively to promote translation, but lacks these features needed to promote translation of the correct pool of mRNAs. Results from yeast studies also allude to the possibility that disrupting eIF4B activity on long structured mRNAs frees ribosomes to translate pro-growth ribosomal protein and metabolic genes with shorter, less structured 5’UTRs. Such observations and those of the present study provoke a reconsideration of eIF4B and its potential role in malignant translation.

Our data present an intriguing notion that eIF4B may play additional roles in mammalian disease beyond its support of eIF4A/F activity – indeed such uncharted action could track opposite to those observed for other related translation initiation factors (e.g., 4A, 4G, and 4). Such a scenario is suggested by the apparently opposite action of eIF4B and eIF4A in PD-L1 induction. Thus, it possible that the translational program active under eIF4B high conditions encourages expression of factors disadvantageous for metastatic growth (e.g., those promoting differentiation vs. stemness, metabolic rigidity vs flexibility, or immunogenicity vs. immune evasion). The influences of eIF4B levels on the process of translation, the specificity of disease-related mRNA pools, or effects on the translation factor interactome may underwrite the changes in metastatic behavior we have observed. Additional study of this initiation factor should elucidate these points as well as the unique relationship between eIF4B and better outcomes for patients with certain breast and other cancers(8).

There is growing appreciation for how mechanisms of immune evasion capable of promoting metastatic growth are controlled at the level of translation (80). The upregulation of inhibitory costimulatory signaling in response to inflammatory cytokines (66) and the integrated stress response (ISR) (81) are prime examples. Altered eIF4E levels also impact the immune infiltration of breast tumors (82). Here we uncover an additional player that negatively impacts the upregulation of PD-L1 in response to the tumoricidal, proinflammatory cytokine IFNgamma. It is interesting that eIF4B knockdown was associated with this and several common indications of tumor-associated immune dysfunction, in both primary (mammary fat pad) and metastatic (i.v. injected) tumors given the distinct effects on disease progression in these models. This could reflect differences in the relative importance of immune evasion versus cellular growth/survival mechanisms impacted by eIF4B. Indeed, there is evidence in the literature suggesting that the ability to upregulate immune suppression via PD-1:PD-L1 activity may be particularly important for metastatic tumor growth and less so for the primary tumor. PD-L1 expression by circulated tumor cells has been reported (83) and may be associated with worse patient outcomes in several cancers (84–86). Comparisons of paired primary and metastatic tumors have revealed disparate frequencies of PD-L1+ tumor cells and unlike immune cells, PD-L1 levels tend to be higher in the primary tumors with PD-L1 protein levels on tumor cells relatively low in the metastatic site and differ across tissues (87, 88). These observations suggest perhaps that the ability to upregulate this inhibitory pathway and the protection it affords may be a limiting in sites of distant tumor colonization. Furthermore, PD-L1 positivity at the metastatic site has been liked to several clinicopathological indicators of poor patient outcome including high Ki-67 index and Lymph node, and metastasis (TNM) stage (89, 90).

Limitations of our study include its largely observational nature. Indeed, further study is needed to shed light on the precise molecular events that underly the novel effects of eIF4B on metastatic growth in vivo and immune evasion. While we demonstrate that modulating eIF4B has significant consequences for these processes, it remains to be seen which transcript pools made into proteins under high- and low-eIF4B conditions account for these effects. Among these, novel regulators in control of pro-metastatic processes, perhaps active only at the level of translation, may be uncovered. Additionally, the impact that abundant or scarce eIF4B may have on the formation of key molecular interactions amongst the translation machinery remains to be investigated (e.g., those between specific eIF4 factors or mRNA). Also, effects on the post-translational modifications that can dictate their activity remains to clarified.

As a technical point, it may be pointed out that our shRNA targeting approach resulted in only partial silencing (knockdown) of eIF4B in these studies, particularly in the EMT6 line that displayed robust baseline eIF4B levels (**Fig. S3A**). However, given eIF4B’s well-recognized role in translation and cell growth, this limitation likely allowed us to model tumor/malignant cell behavior associated with eIF4B levels at the lower end of the expression spectra without the expected detrimental effects likely to arise from complete loss. It is important to note that despite drawbacks of this approach, we were able to generate a nuanced picture of how eIF4B levels impact the in vivo disease course of breast cancer – one that at least superficially resembles the relationship between this factor and patient outcomes according to publicly deposited survival data. Additional studies are needed to confirm whether the survival differences seen with divergent eIF4B levels stem from differential immune evasion or additional mechanisms.

Nevertheless, our findings may have significant implications for the prognosis and treatment of breast cancers. For example, our results suggest that tumors with low eIF4B levels may be more likely to progress to metastatic disease and thrive. The ability to predict which tumors are likely to reach this life-threatening stage would greatly aid caregivers in critical treatment decisions such as when to apply or withhold aggressive cytotoxic therapies. Additionally, while checkpoint blockade immunotherapies have revolutionized the treatment of many cancers, reliable identification of patients likely to respond to agents targeting the PD-1:PD-L1 axis remains critical for both clinical studies and treatment. Our study suggests that eIF4B levels, and perhaps more broadly, the relative balance between initiation factor levels, may be useful in identifying patients with robust capacity for immune suppression that may nevertheless be responsive to PD-1 or PD-L1 targeting.

## Materials and Methods

### Cell lines and mice

The murine TNBC cell line 4T1 was obtained from ATCC (Rockville, MD). EMT6 cells as well as derivative sub-lines of both 4T1 and EMT6 lines were kind gifts of Dr. Craig Brackett (Roswell Park). Luciferase-expressing EMT6 cells were generated by transfecting the parental line with a plasmid DNA encoding a CMV promoter-driven firefly luciferase (Luciferase-pcDNA, prepared in X420 as pcDNA3_CMV-Fluc; Addgene®, cat.# 18964) using Polyplus® JetPrime™ as per the manufacturer’s protocol. Transfectants were grown under selection (0.75 mg/ml G418, VWR®) to establish stable derivative lines. 4T1-luc cells were provided by Dr. John Ebos (Roswell Park). All cell lines were maintained as adherent cultures in “complete” RPMI media (cRPMI, Gibco) supplemented with 10% fetal bovine serum (Gemini Bio), 1% penicillin/streptomycin, and 1% L-Glutamine (cRPMI) and grown at 37°C in a 5% CO2. Stable shRNA/control vector-expressing 4T1 and EMT6 lines were derived from both parental and luciferase-expressing lines.

Wild type female BALB/c mice aged 6-8 weeks were purchased from Jackson Laboratories. Rag2^−/−^ γc^−/−^ (C;129S4-*Rag2^tm1.1Flv^ Il2rg^tm1.1Flv^*/J Strain #:014593) founders were similarly obtained and bred in house. All mice were housed under specific pathogen free conditions in Roswell Park’s Laboratory Animal Shared Resource, and all procedures involving live animals were approved by the IACUC and carried out in accordance with institutional and federal guidelines ensuring animal welfare.

### Immunoblot analysis

Cell pellets were collected, lysed with RIPPA buffer then samples were normalized based on protein levels quantified by BCA assay. Alternatively, pellets of known cell numbers were collected, resuspended in Laemmli Buffer (BioRad), and briefly boiled before clarification by centrifugation. Samples were resolved by SDS/PAGE before transfer to nitrocellulose membranes via Turboblot (BioRad) for probing with primary and secondary antibodies (Abcam). Blots were visualized using a LiCor Odyssey Imager. Resulting band densities were quantified and normalized to loading controls by Image J software.

### Lentiviral transduction and generation of stable shRNA expressing cell lines

shRNA constructs designed to target murine eIF4B (clone IDs: RMM3981-201802392, RMM3981-201807177, RMM3981-201811768) were purchased from Dharmacon, Inc. Glycerol stocks of bacteria carrying these and a pLKO.1 eGFP positive control vector were propagated, and plasmid DNA was isolated using OMEGA’s EZNA plasmid DNA miniprep kit (Omega Biotek Inc Cat# D694202). Lentivirus containing supernatants were produced in packaging (293T) cells transfected with individual elF4B shRNAs or the pLKO.1 eGFP positive control using LipoD293™ transfection reagent (SignaGen Laboratories; Cat# SL100668) at a 1:2.5 DNA:Lipo ratio (2.5ug transfer plasmid + 2.5ug psPAX2 + 1.0ug pMD2.G and 15ul lipoD per plate). 48 and 72 hrs post-transfection, supernatants were harvested, filtered, aliquoted, and stored at −80°C. For transduction, viral supernatants were diluted 1:3 in growth media containing 4ug/ml polybrene, added to a culture plate seeded with EMT6 or 4T1 cells which was placed in a microtiter rotor and spun at 1800 rpm for 45 minutes at room temperature. Stable transductants were expanded and selected for by culture in the presence of puromycin (2ug/ml).

### In vitro assays of tumor cell growth

The impact of eIF4B levels on breast cancer growth rate was assessed by seeding the indicated lines in triplicate cell culture plates followed by periodic detachment and manual cell counting. Effects on growth and in vitro migration on cell culture-treated plastic were observed using a scratch assay. Here, breast cancer cell lines were grown to near confluency in 6-well culture plates. A wound/scratch was made using a p-200 pipette tip, and debris was gently washed away. Brightfield images of the scratch area were taken at 0h, 10h, 15h, and 24h later using an inverted microscope mounted camera. Between time points, the plates were incubated at 37°C, 5% CO2. Quantification of the gap area and its change over time was achieved using ImageJ software.

The capacity for invasive growth was also assessed by an in vitro Extracellular matrix (ECM) invasion assay. Here, transwell inserts compatible with fluorescence microscopy (8um pore size) were pre-coated with ECM (Geltrex, Life Technologies) as described elsewhere(91), and placed into 24-well plate wells containing either serum-free RPMI or media supplemented with 10% serum. Cell lines were harvested from sub-confluent growth flasks and pre-stained with 10uM cell trace violet (Molecular probes). After quenching, and washing the cells, they were resuspended in serum-free RPMI and their concentrations were adjusted (5×10^4^ cells/100ul). These were carefully added in triplicate into the top chambers of the transwell plates, and fluorescent images of the underside of the transwell were collected over a 24hr period using a Cytation multi-mode automated digital imager (Biotek). Data were presented as mean fluorescence intensity (MFI) over 5-9, 5X fields/well.

### Puromycin incorporation (SUnSET) assay

Non-radiological measurement of global translation was carried out by SUnSET assay (57). Here, cell lines were plated and grown to 50-60% confluency. 0.25-0.5 ×10^6^ cells were transferred to 96 well plate and pulsed with puromycin (10μg/ml in cRPMI) for 15 minutes at 37°C, 5% CO2. After replacing the media with fresh media (without puromycin) and an additional incubation for 35 min., surface puromycin was detected by immunostaining with anti-puromycin monoclonal antibody (clone 12D10; Millipore-Sigma) followed by flow cytometry analysis.

### In vivo tumor model experiments

For tumor challenge experiments, the indicated luciferase-expressing cell lines were grown to near confluency, detached with trypsin/EDTA, washed, and resuspended in sterile PBS. For intra-mammary (i.m.) tumor challenge, 5×10^4^ tumor cells were implanted into the fourth mammary fat pad of hand restrained Balb/c mice (n = 5-8/group). Primary tumor dimensions were measured every 2-3 days by digital caliper, and volumes were determined according to the formula V=½(Length)xWidth^2^. 21-28 days post-implantation mice were euthanized (prior to reaching 2cm in the longest dimension). Tumor-infiltrating leukocytes (TILs) were recovered by enzymatic digestion. Tumor-draining and tumor-distal lymph nodes as well as spleen cell suspension were also generated by mechanical dissociation prior to flow cytometric characterization. For studies of pulmonary metastases, 5.0×10^4^ tumor cells were injected intravenously (i.v.) by the tail vein (n = 5/group), and mice were monitored daily for signs of distress. Surrogate survival times were determined based on the need for humane euthanasia.

In the above studies, primary and disseminated tumor burden was also observed by periodic, luciferase-dependent bioluminescence imaging (BLI) using Xenogen IVIS Spectrum technology (Perkin-Elmer, Waltham, MA). On the indicated day post-implantation, tumor-bearing mice were anesthetized by isoflurane inhalation and d-luciferin (Gold Biotechnology; 150mg/kg mouse weight) was administered by intraperitoneal (i.p.) injection. Mice were immediately placed in the warmed imaging chamber and anesthesia was maintained during the imaging session. In some experiments, secondary tumor (metastatic) growth was visualized by covering the primary tumor with imaging dampeners. Photonic flux (photon per second) in representative images was quantified with Living Image software (V4.7.3; Perkin-Elmer).

### Flow cytometry Analysis

Single cell suspensions of tumor infiltrating leukocytes (TILs) and cells from tumor associated lymphoid tissues were recovered from mice implanted with the indicated TNBC lines. Cells were washed with phosphate buffered saline (PBS) supplemented with 2% v/v fetal bovine serum (FBS; Gemini Bioproducts) and staining with panels of fluorochrome-conjugated antibodies specific for immune cell type-defining markers. Intracellular markers (e.g., FOXP3, KI67) were also stained after fixation and permeabilized using a specialized kit (ebiosciences) according to the manufacturer’s specifications. Multi-color flow cytometry data was collected using a BD LSR II analyzer and analyzed using FlowJo software (v10.7, BD Biosciences). For a complete list of antibodies used in this study, see **Supplemental Table S1**.

### RT-PCR analysis

For transcript measurements by RT-PCR, total RNA was recovered from harvested cells and tissues using Trizol (Thermo Scientific) and purified by chloroform and isopropanol precipitation. cDNA was generated using the high-capacity cDNA synthesis kit (Applied Biosystems) as per the manufacturer’s specifications with 1ug purified RNA. qRT-PCR analysis was carried out using PowerUp™ SYBR™ Green Master Mix (Applied Biosystems), validated primer pairs obtained from IDT based on sequences obtained from Primer Bank (https://pga.mgh.harvard.edu/primerbank/index.html. For all experiments, gene expression levels were normalized to the housekeeping gene RPL13a expression level and compared to the levels of EMT6/4T1 sh-sc control. All qRT-PCR reactions were performed in technical triplicate with samples from at least 3 independent experiments using a Bio-Rad CFX96 thermal-cycler. Fold change was calculated from gene threshold (Cq) values using the equation (2^−∆∆Cq^) and assessed in Microsoft Excel. A complete list of primers used in this study can be found in **Supplemental Table S2**.

### Leukocyte-tumor co-culture assay

Tumor cells were plated at a density of 15,000 cells per well in a 24-well plate in cRPMI. The splenocytes from a wild type female Balb/c mouse (aged 6-8 weeks) were excised, dissociated, and depleted of RBC by a short (~30 second) incubation in ACK lysis buffer (Gibco). Cells were washed and resuspended before counting and stained with a 10 μM working solution of Cell Trace Violet (Molecular Probes) as per the manufacturer’s recommendations. Labelled splenocyte cells were then added to tumor cell-coated wells (1×10^6^/1.5ml/well) in cRPMI) with and without stimulating anti-CD3/CD28 antibodies (both at a final concentration of 1ug/ml). 72 hr later, non-adherent and adherent cells were collected by rinsing and brief trypsinization. Surface and internal phenotypic markers were then stained for flow cytometry analysis.

### In vitro PD-L1 induction

The indicated cell lines were seeded in 6-well plates and allowed to reach 60-70% confluency. Cells were then treated with recombinant IFNγ (Peprotech; 100 ug/ml) or media alone as in prior studies (66). After incubation at 37°C 5%CO2 for 24h, cells were mechanically harvested, and surface PD-L1 was probed for with fluorescently conjugated anti-PD-L1 antibody. Baseline (untreated cells) and IFNγ-induced PD-L1 levels were then determined by flow cytometry, and corroborative measurements were made using RT-PCR.

### Analysis of Breast Cancer Patient Survival and Gene Expression Data

Using the Human Protein Atlas (44) and KM Plotter analysis tool for RNA-seq data (92), the relationship between eIF4B transcript and survival of Breast Invasive Carcinoma (BRCA) (n = 1075) patients was examined using Cox proportional hazards regression and visualized by Kaplan-Meier plots. Similar analyses were carried out using KM Plotter (93) and published data on eIF4B protein levels and patient survival from a prior study (68). Compiled survival and gene expression (microarray) data from at least 33 breast cancer datasets were also analyzed using KM plotter (94). Here, biased arrays and redundant samples were excluded, and optimal probe sets were selected based on specificity, coverage, and degradation resistance by Jetset (95). In these studies, patients in each dataset were stratified by eIF4B expression (above and below median value) and compared based on overall survival (OS), progression free survival (PFS), distant metastasis free survival (DMFS), or post-progression survival (PPS). Hazard Ratio (HR), 5-year survival rates and upper quartile survival times were found. An HR>1 indicates a survival disadvantage. To compare eIF4 family member expression levels across normal and cancerous breast tissue as well as various subcategories of breast cancers and tumor characteristics, RNASeq data or microarray (U133 Plus 2.0) data was analyzed using the Human Protein Atlas or the GENT2 analysis tool (43). Relative eIF4B mRNA levels across human breast cancer lines in the Cancer Cell Line Encyclopedia (CCLE) (49) were found using cBioPortal.org and were represented by a heat map generated with Graphpad Prism.

### Statistical Analysis

Significant differences between two groups were determined by standard t test, unless otherwise indicated. Quantified data are presented as means with error bars presenting SEM. Survival (Kaplan Meier) curve analysis for mouse experiments was performed using Graphpad Prism (v8) software or the indicated web-based analysis tool (human datasets).

## Acknowledgments

This work was supported by NIH grants R00GM119173 and R01GM139977 (S.E.W.), and institutional funds from the University at Buffalo College of Arts and Sciences (S.E.W.) and Roswell Park Comprehensive Cancer Center (J.B.). This work involved the use of the Shared Resources at Roswell Park supported by National Cancer Institute (NCI) grant P30-CA016056. The authors would like to thank Dr. Santosh Patnaik and Eric Kannisto (Roswell Park) for helpful discussion and generation of a luciferase expressing EMT6 cell line.

## Supplementary Figure Legends

**Supplementary Figure S1.**
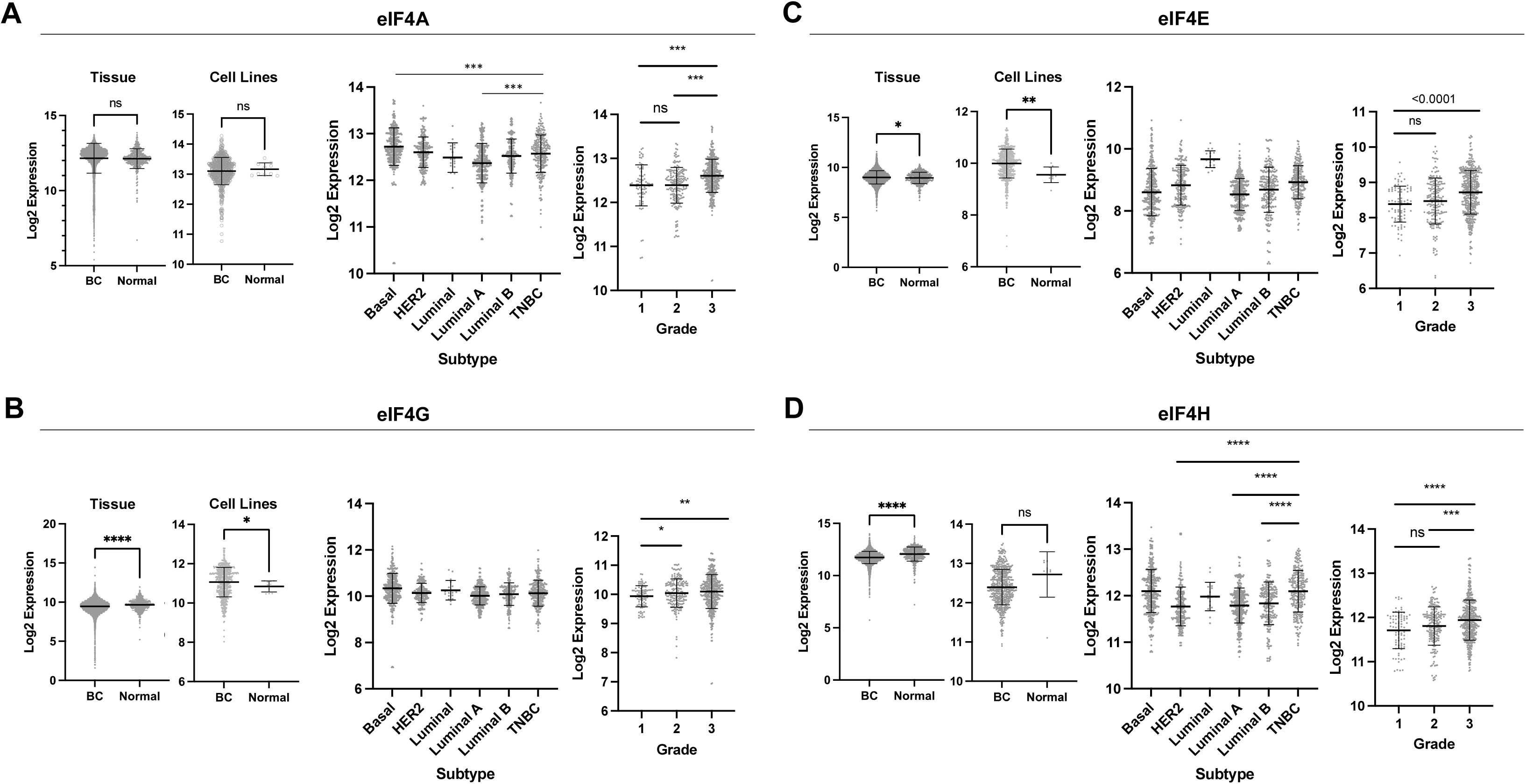
Expression of translation initiation factors across breast cancer subtypes and grades. Gene expression (GeneChip microarray) datasets were accessed through the GENT2 analysis webtool, and the levels of eIF4A **(A)**, eIF4G **(B)**, eIF4E **(C)**, and eIF4H **(D)** transcript were assessed across normal breast and breast cancer tissue samples (left) and tumors of different breast cancer subtypes (middle) and grades (right). Shown are the mean values (bars) for each group +/-SEM. Statistical differences (by unpaired t test) are shown. (ns = not significant, \**p* < 0.05, **p<0.01, ***p<0.001).

**Supplementary Figure S2.**
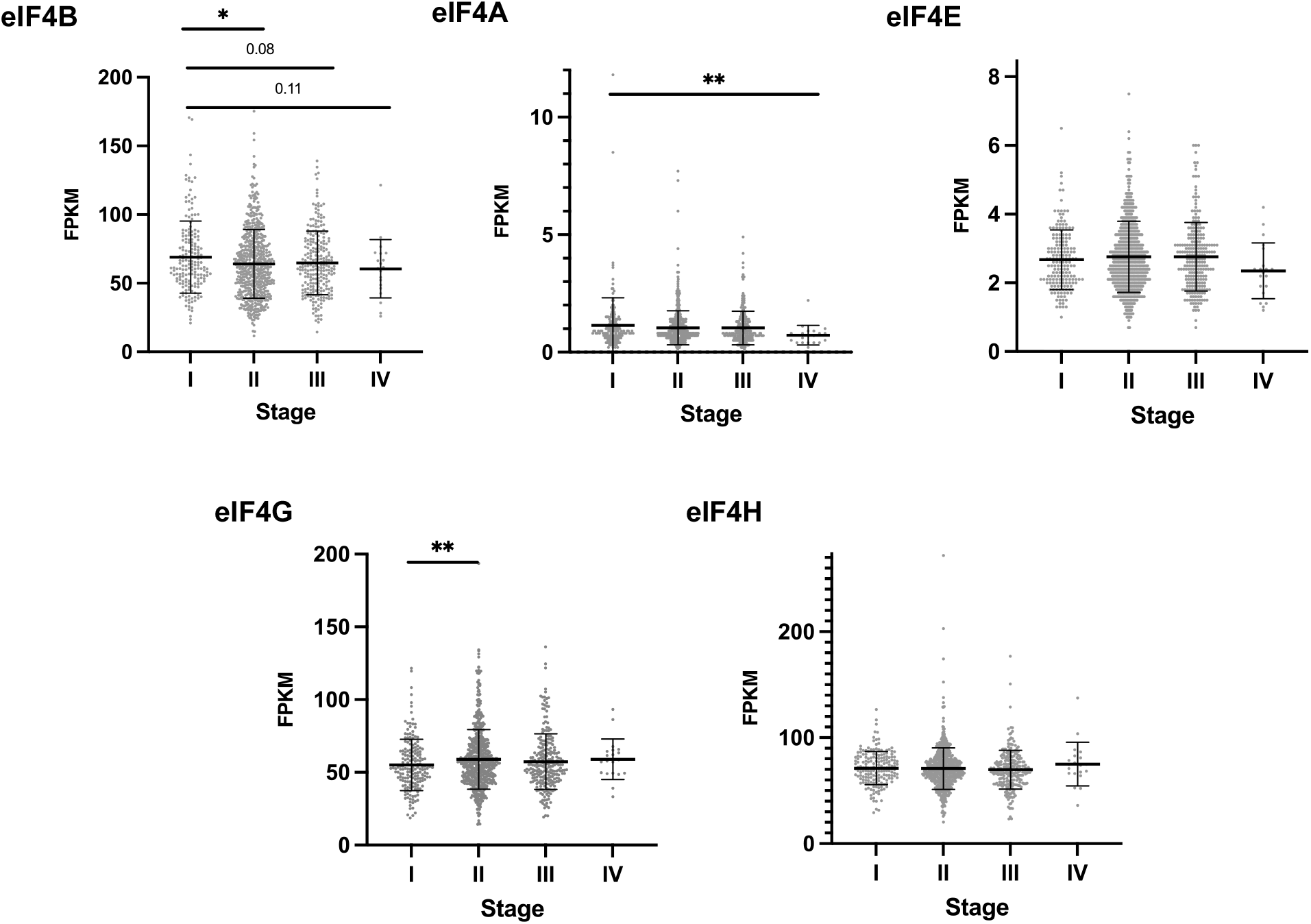
Expression of translation initiation factors across breast cancer stages. RNASeq data available for Breast Invasive Carcinoma (BRCA) patients in the TCGA Cancer Genome Atlas (TCGA) was used to establish initiation factor expression patterns across stage I, II, III, and IV breast tumors. Shown are the mean Fragments per kilobase of transcript per million mapped fragment (FPKM) values (bars) for each group +/-SEM. Statistically significant differences between stages are indicated (\**p* < 0.05, **p<0.01, by unpaired t test).

**Supplementary Figure S3.**
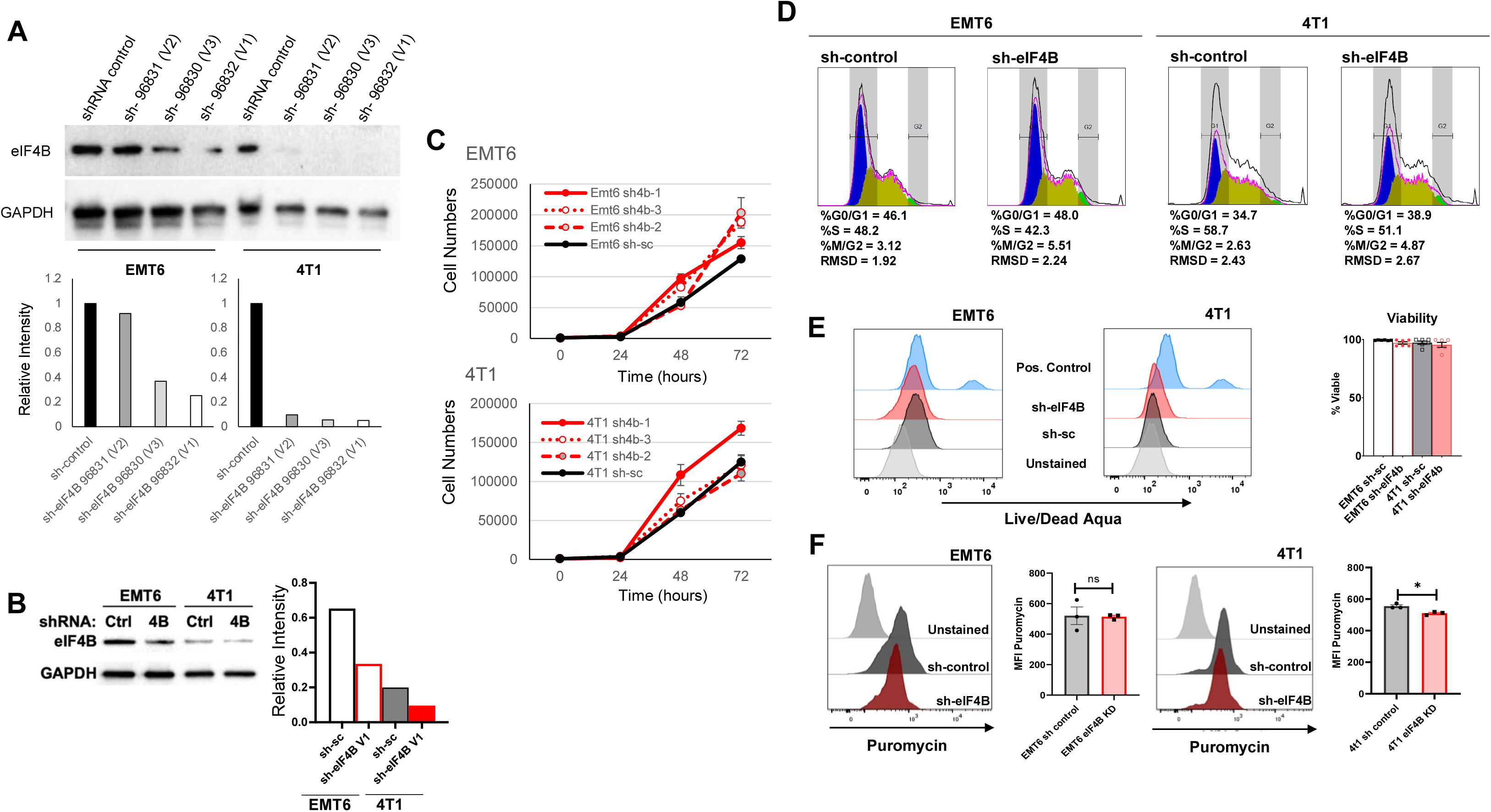
shRNA-mediated eIF4B knockdown and its effects on growth, cell cycle, viability, and global translation in vitro. shRNA constructs targeting eIF4B expression or inert control constructs were delivered to EMT6 and 4T1 cell lines by lentiviral transduction. Stable transductant lines were generated, and **(A)** the ability of three separate shRNA knockdown constructs to reduce eIF4B protein levels in both cell lines was confirmed by Western Blot with a GAPDH loading control and normalized to relevant sh-control cells. **(B)** Similarly, sh-eIF4B and control lines were generated by transduction of luciferase reporter-expressing EMT6 and 4T1 cell lines used for in vivo experiments. **(C)** The in vitro growth of stably transduced lines carrying eIF4B targeting shRNA or control constructs was observed by counting cell numbers in triplicate seeded culture plates over time. **(D)** The distribution of cells across cell cycle stages as indicated by propidium idodine uptake was assessed by flow cytometry using Flowjo software. **(E)** The viability of the aforementioned transductants grown in standard culture conditions was determined based on exclusion of Live/Dead Aqua stain measured by flow cytometry. **(F)** Effects of eIF4B knockdown on global translation efficiency in murine breast cancer lines was assessed in a puromycin incorporation SUnSET assay. Shown are representative findings from 2-3 independent experiments with samples run in technical triplicate (panels C,B, E). Error bars depict the SEM. *p<0.05 by t test.

**Supplementary Figure S4.**
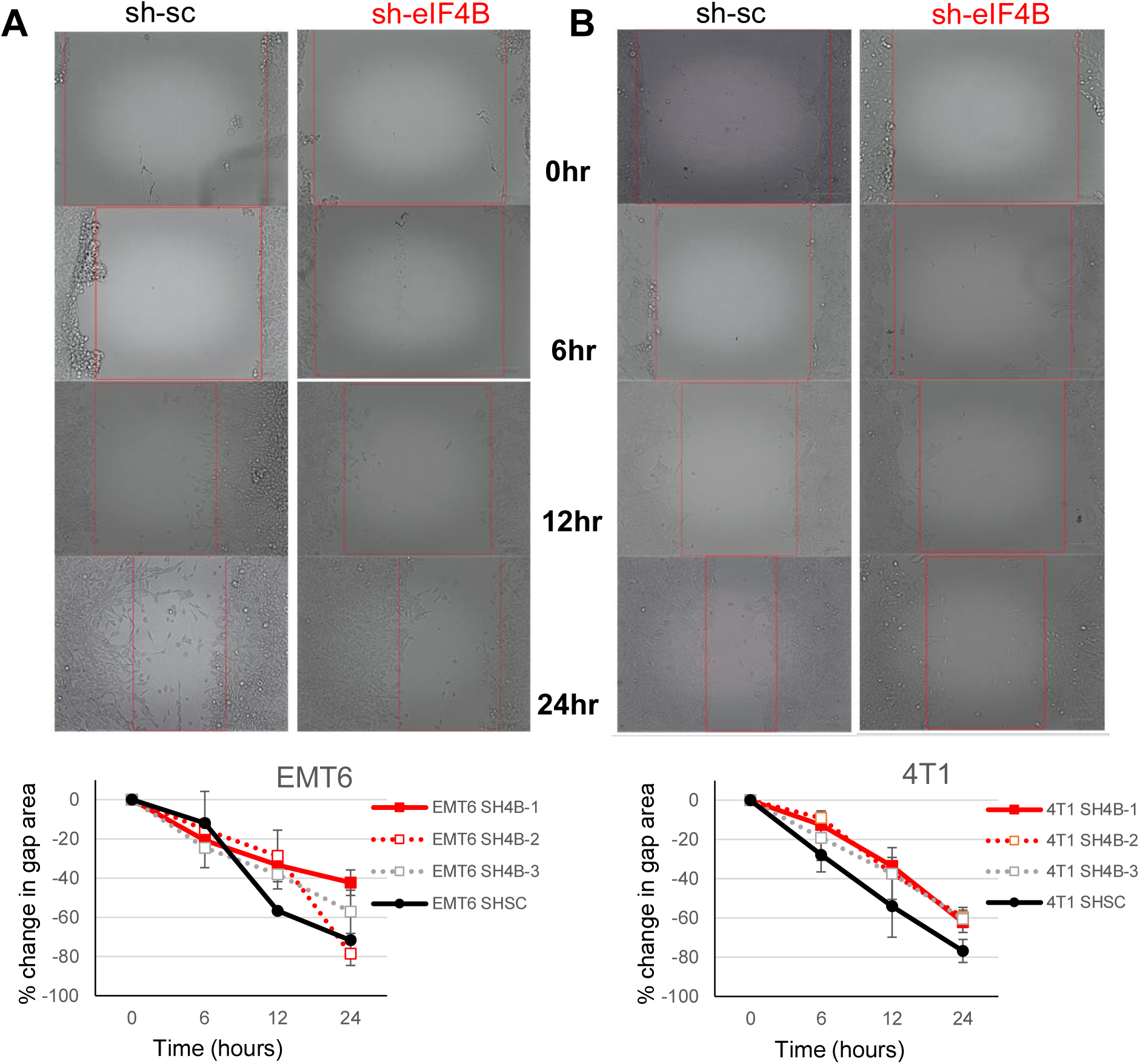
Effects of eIF4B knockdown on in vitro growth and migration. Stable transductants carrying either eIF4B targeting shRNA or control constructs were seeded in 6-well culture plates and grown to confluency. A scratch in the monolayer was inflicted, and the area of the “wound” was measured overtime. The cell-free gap for each triplicate sample was photographed periodically and measured using ImageJ software. Shown are representative micrographs and the mean change in gap area over time (+/-SEM) from one of three independent experiments.

**Supplementary Figure S5.**
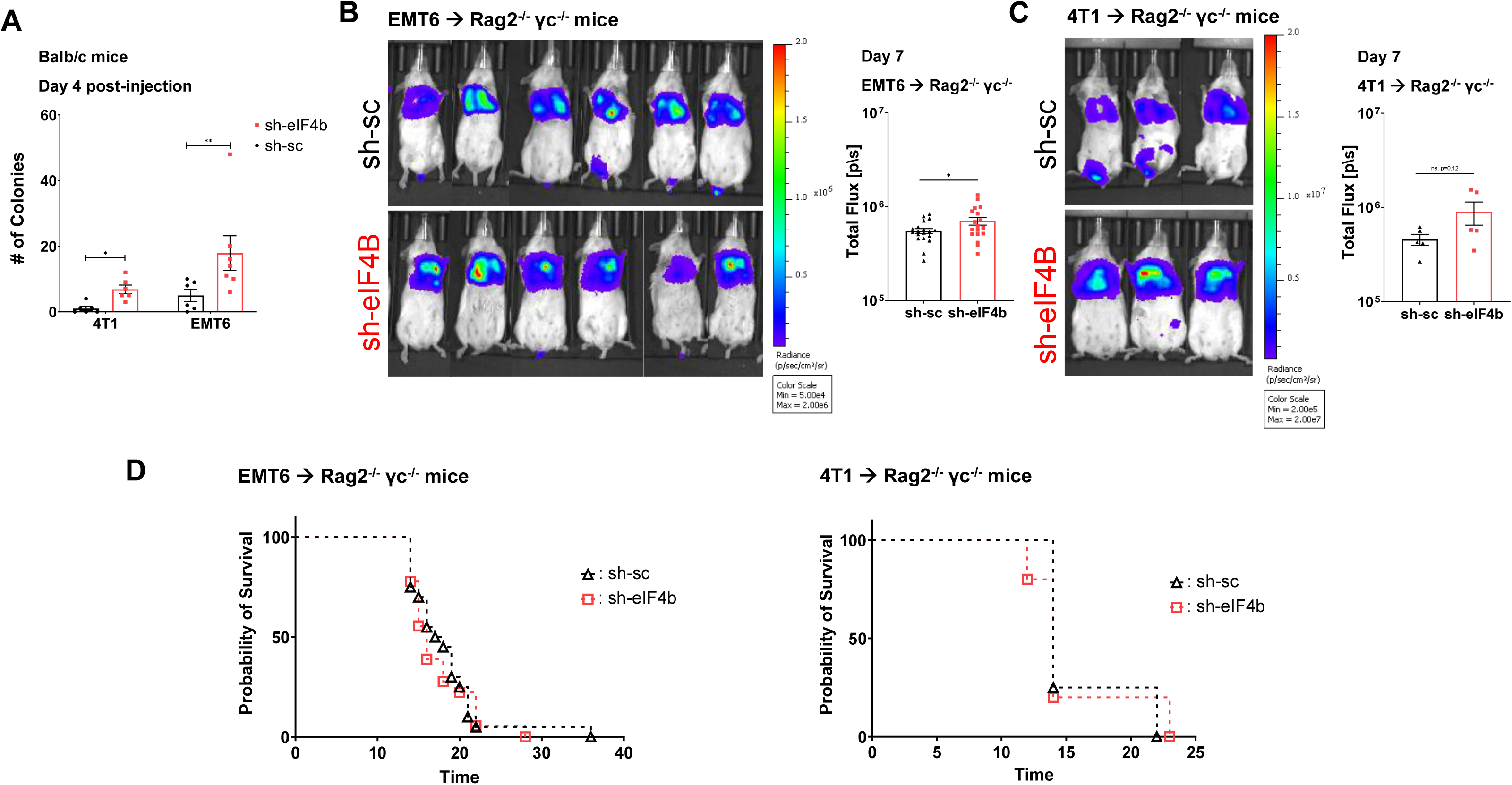
Consequences of eIF4B knockdown on metastatic tumor development in normal and immunodeficient mice. Luciferase expressing breast cancer cell lines with experimentally modulated eIF4B levels (and corresponding control lines) were injected i.v. into wild type Balb/c mice (panel **A)** and syngeneic double knockout mice with simultaneous genetic deletion of Rag2 and IL2receptor gamma chain expression (Rag2-/-,γc-/-; panels **B-D**). **(A)** The establishment of nascent metastases in cohorts of Balb/c mice (n= 3-4mice/experiment) was observed within four days of challenge. Here lung tissue homogenates were plated ex vivo for 10-14 days, and adherent tumor cell colonies were visually quantified after fixation and crystal violet staining. Metastatic tumor burden in Rag2-/-,γc-/-mice (n=5-8 mice/trial) was monitored weekly by IVIS after i.v. challenge with the indicated EMT6 **(B)** and 4T1 **(C)** cells. Representative IVIS images (left) and mean tumor burdens +/-SEM (right) are shown for the day 7 time point of two independent experiments. **(D)** Survival of these mice after i.v. tumor challenge was determined by Kaplan-Meier analysis. Panel A depicts the mean +/-SEM results of two independent trials. Panels B and C show representative IVIS images (left) and mean tumor burdens +/-SEM (right) and panel D depicts pooled results from two independent experiments. \**p* < 0.05, **p<0.01 by t test.

**Supplementary Figure S6.**
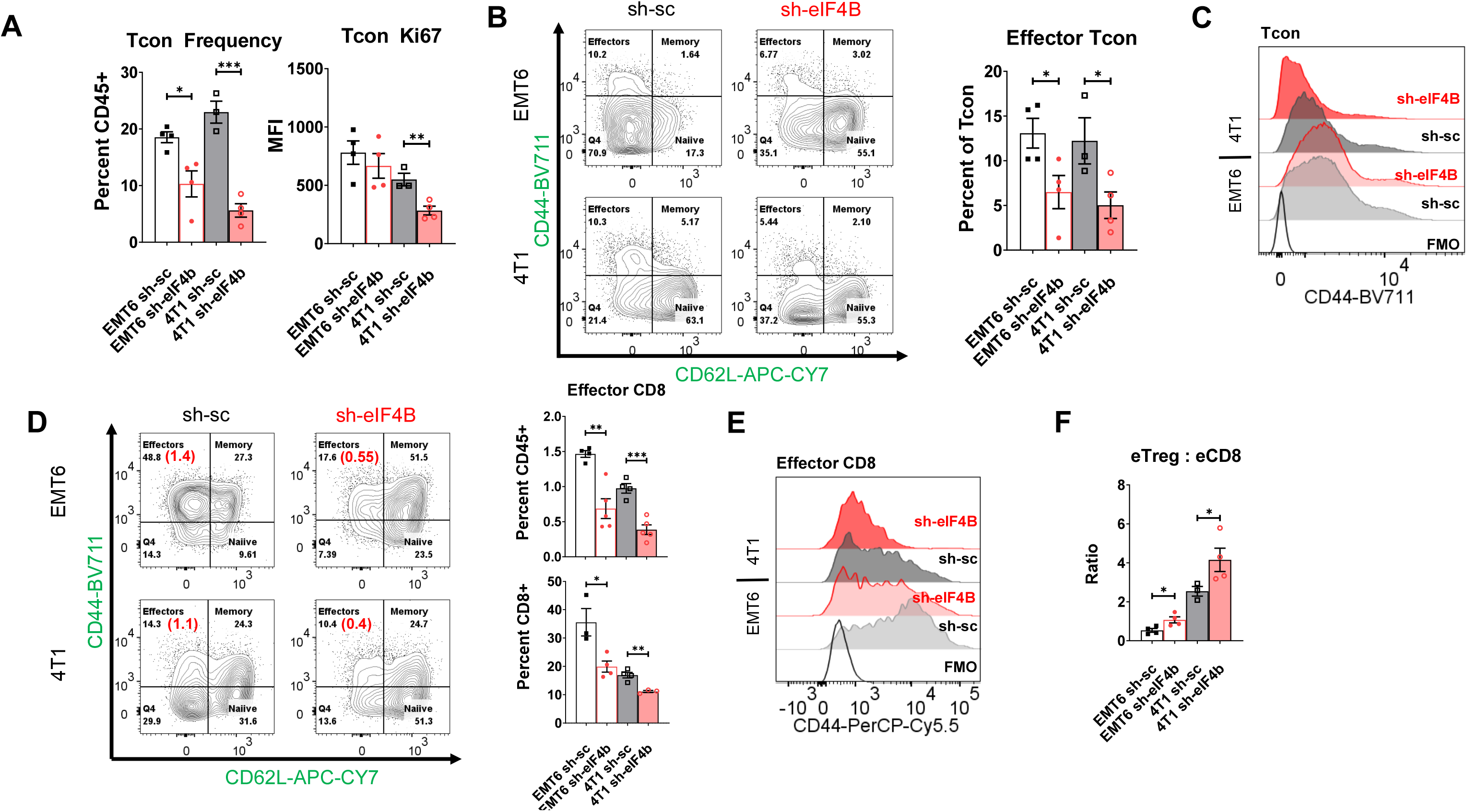
Effects of eIF4B knockdown on the T cell response to pulmonary breast cancer metastases in mice. Balb/c mice were challenged as described for Fig. 3 with the indicated breast cancer cell lines injected i.v. to model metastatic tumor development. Tumor-bearing lungs were excised and digested enzymatically yielding single cell suspensions that were characterized by multi-color flow cytometry. The proportions of non-regulatory (Tcon) CD4+ cells among the immune (CD45+) cells recovered from tumor-bearing lungs and their proliferative capacity based on levels of KI67 **(A)** were thus determined. Surface marker (CD44 and CD62L) profiles identified the activated effector T cells among these cells **(B,C)** as well as potential cytotoxic CD8+ T cells **(D, E)**. Additionally, the abundance of effector CD8+ T cells (eCD8s) relative to Tregs displaying an activated, effector phenotype (eTregs) in these samples (i.e., the eTreg:eCD8 ratio) was also determined **(F)**. Panels depict representative flow cytometry plots and corresponding quantification of key markers and cell types (mean +/-SEM) from three independent experiments. (\**p* < 0.05, **p<0.01) by t test.

**Supplementary Figure S7.**
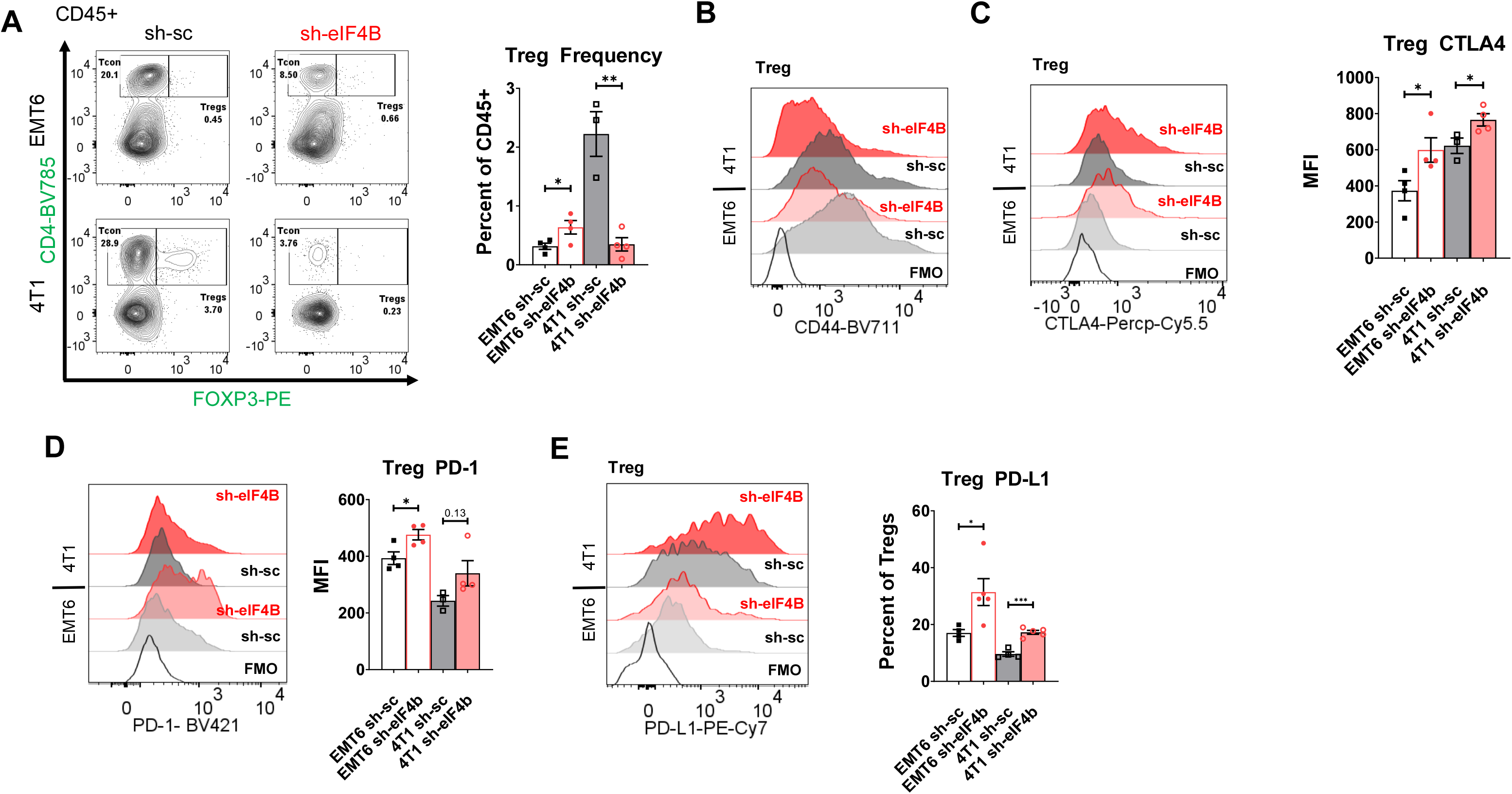
Effects of eIF4B knockdown on the Regulatory T cells in the immune microenvironment of metastatic tumors. Balb/c mice were challenged as described for Fig. 3 with the indicated breast cancer cell lines injected i.v. to model metastatic tumor development. The immune cells recovered from tumor-bearing lungs were characterized by multi-color flow cytometry. The frequencies of Tregs (CD4+/CD25+/Foxp3+) among the lung tumor associated immune (CD45+) cells **(A)** were determined as well as the levels CD44, PD-1, CTLA-4, and PD-L1, **(B-E)** markers of Treg activation and function. Shown are representative findings (mean +/-SEM) from three independent experiments. (\**p* < 0.05, **p<0.01) by t test.

**Supplementary Figure S8.**
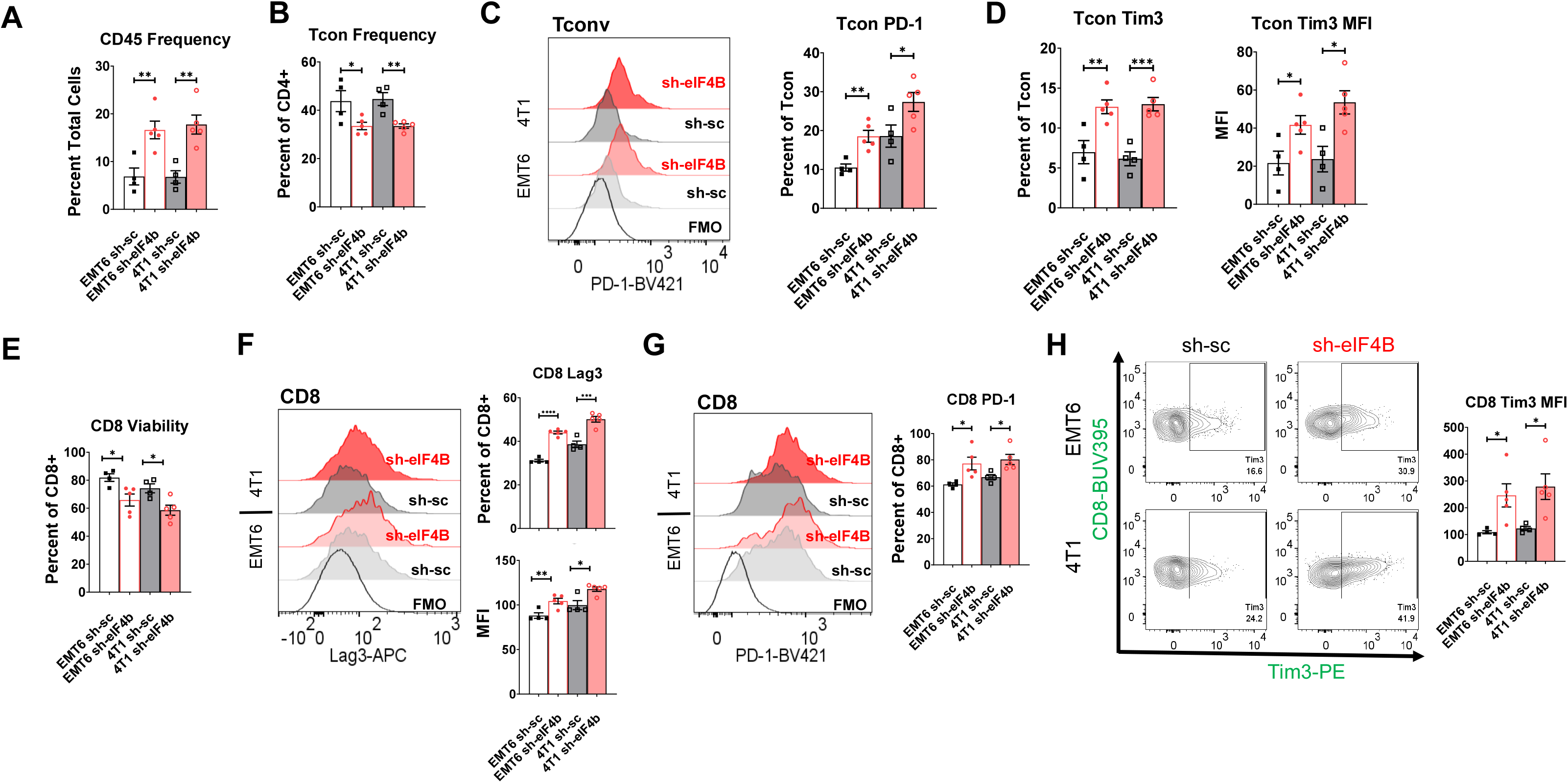
Characterizing the impact of eIF4B knockdown on effector T cells in the primary tumor niche. Balb/c mice were challenged as described for Fig. 3 with the indicated breast cancer cell lines implanted into the mammary fat pad to model primary tumor growth. At the conclusion of the experiment, mammary tumors were dissected and enzymatically digested yielding single cell suspensions that were characterized by multi-color flow cytometry. The degree of immune cell (CD45+) infiltration **(A)** and the relative size of the non-regulatory (Tcon) CD4+ T cell compartment among these tumor-infiltrating leukocytes (TILs) **(B)** were thus determined along with the expression of immune checkpoint molecules PD-1 **(C)** and Tim3 **(D)** among these cells. The viability CD8+ TILs **(E)** and the expression of Lag3, PD-1, and Tim3 **(F-H)** by these cells were similarly found. Shown are representative findings (mean +/-SEM) from three independent experiments. (\**p* < 0.05, **p<0.01, ***p<0.001) by t test.

**Supplementary Figure S9.**
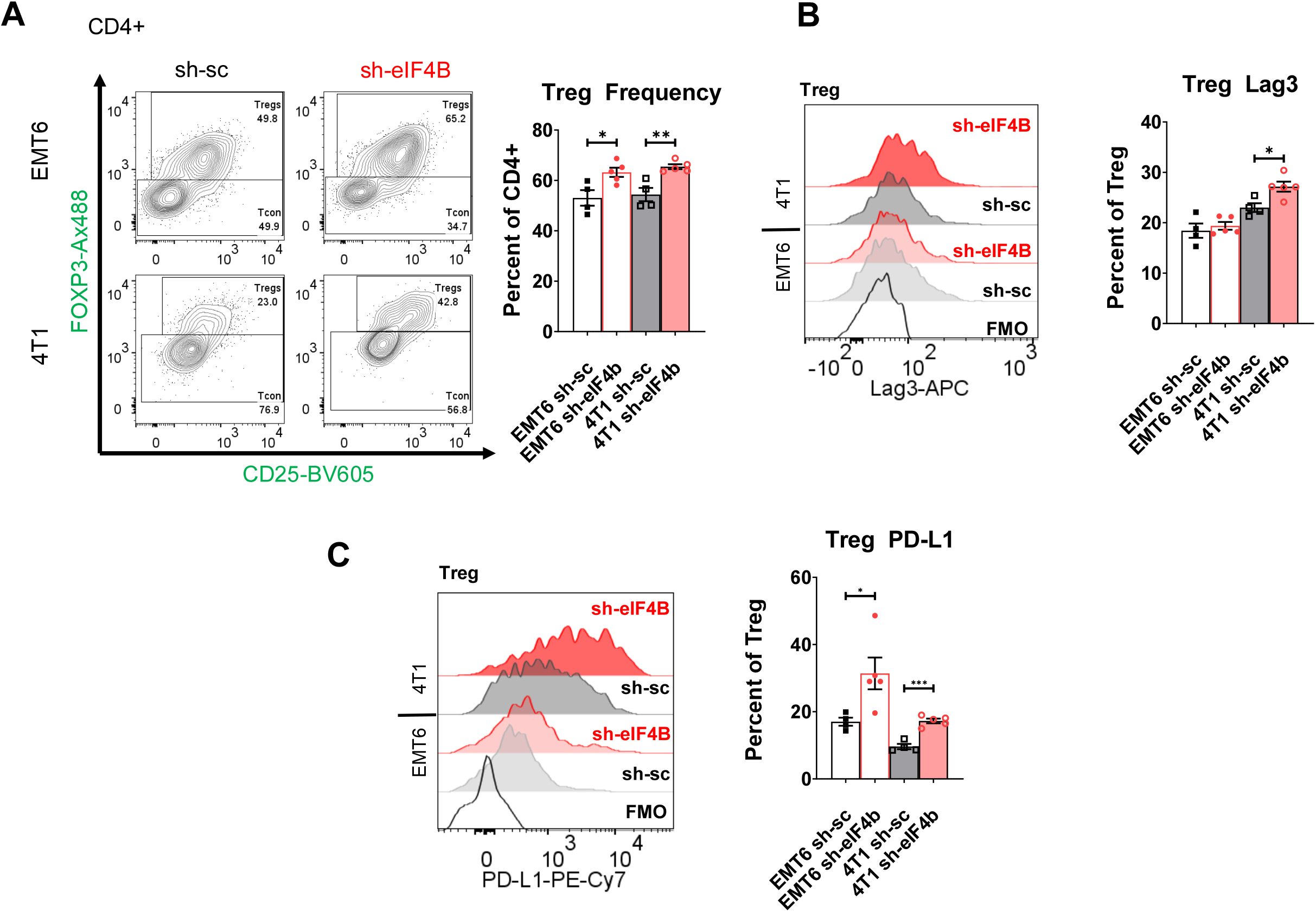
Regulatory T cells in the primary tumors of mice implanted with control and eIF4B knockdown TNBC lines. Balb/c mice were challenged with the indicated breast cancer cell lines implanted into the mammary fat pad to model primary tumor growth as described for Fig.3. At the end of the experiment, mammary tumors were dissected and enzymatically digested yielding single cell suspensions that were characterized by multi-color flow cytometry. The frequency of Treg (Foxp3+/CD25+) cells among intra-tumoral CD4+ T cells were thus determined for all groups **(A)** as well as the prevalence of immune checkpoint factor Lag3 **(B)** and PD-L1 **(C)** expression by these suppressor cells in the tumor niche. Shown are representative findings (mean +/-SEM) from 3 independent experiments. (\**p* < 0.05, **p<0.01) by t test.

**Supplementary Figure S10.**
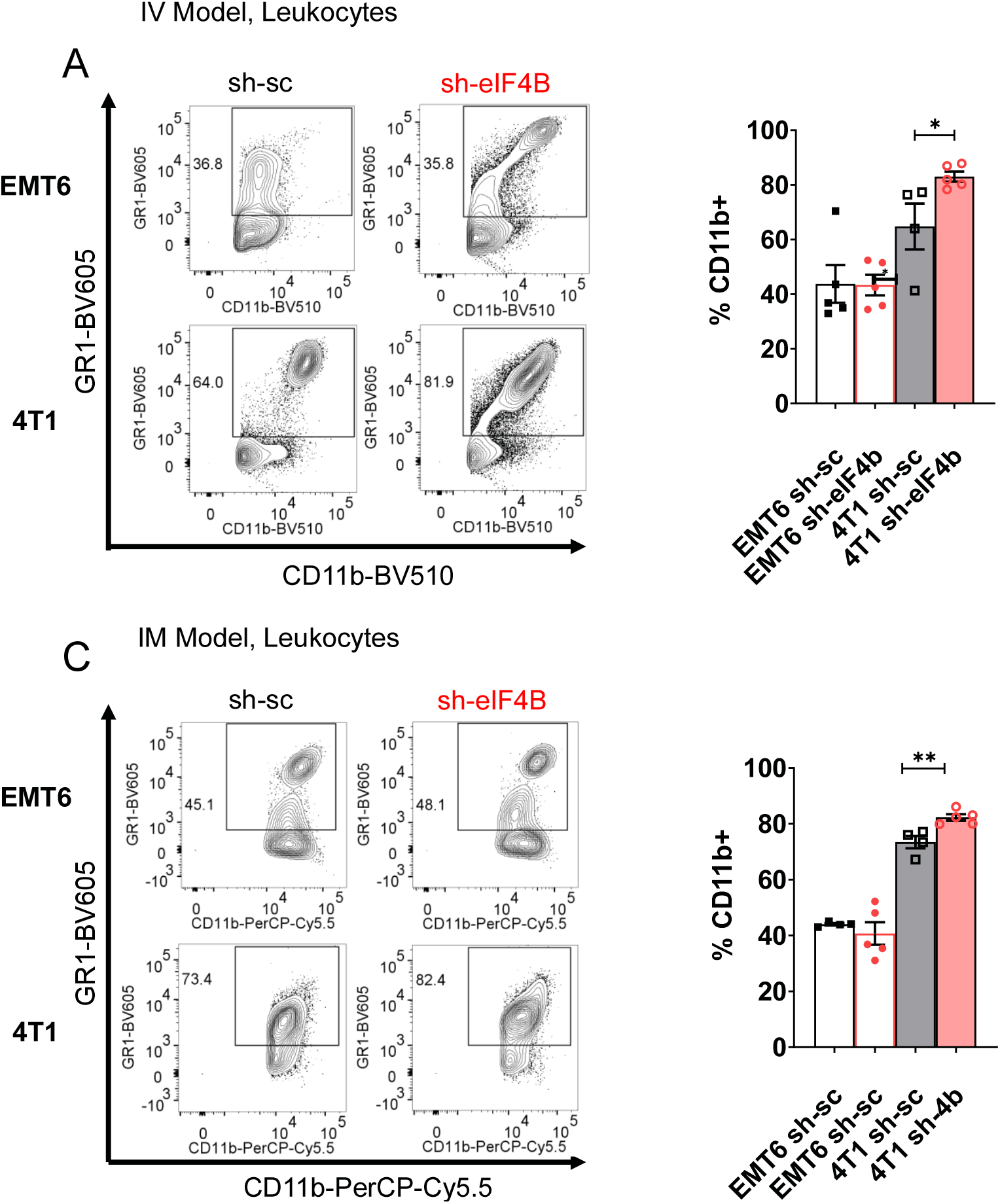
Effects of sh-RNA mediated reduction of eIF4B on myeloid derived suppressor cell frequencies across tumor models. Balb/c mice were challenged with the indicated breast cancer cell lines in models of metastatic tumor development (i.v. injection) and primary tumor growth (i.m. implantation) as described for Fig. 3. Tumor-bearing lungs **(A)** and excised mammary tumors **(B)** were enzymatically digested yielding single cell suspensions that were characterized by multi-color flow cytometry. The frequencies of cells displaying a surface marker profile suggestive of myeloid derived suppressor cells (CD11b+/GR1+) were found. Shown are representative findings (mean +/-SEM) from 2-3 independent experiments. (\**p* < 0.05, **p<0.01) by t test.

**Supplementary Figure S11.**
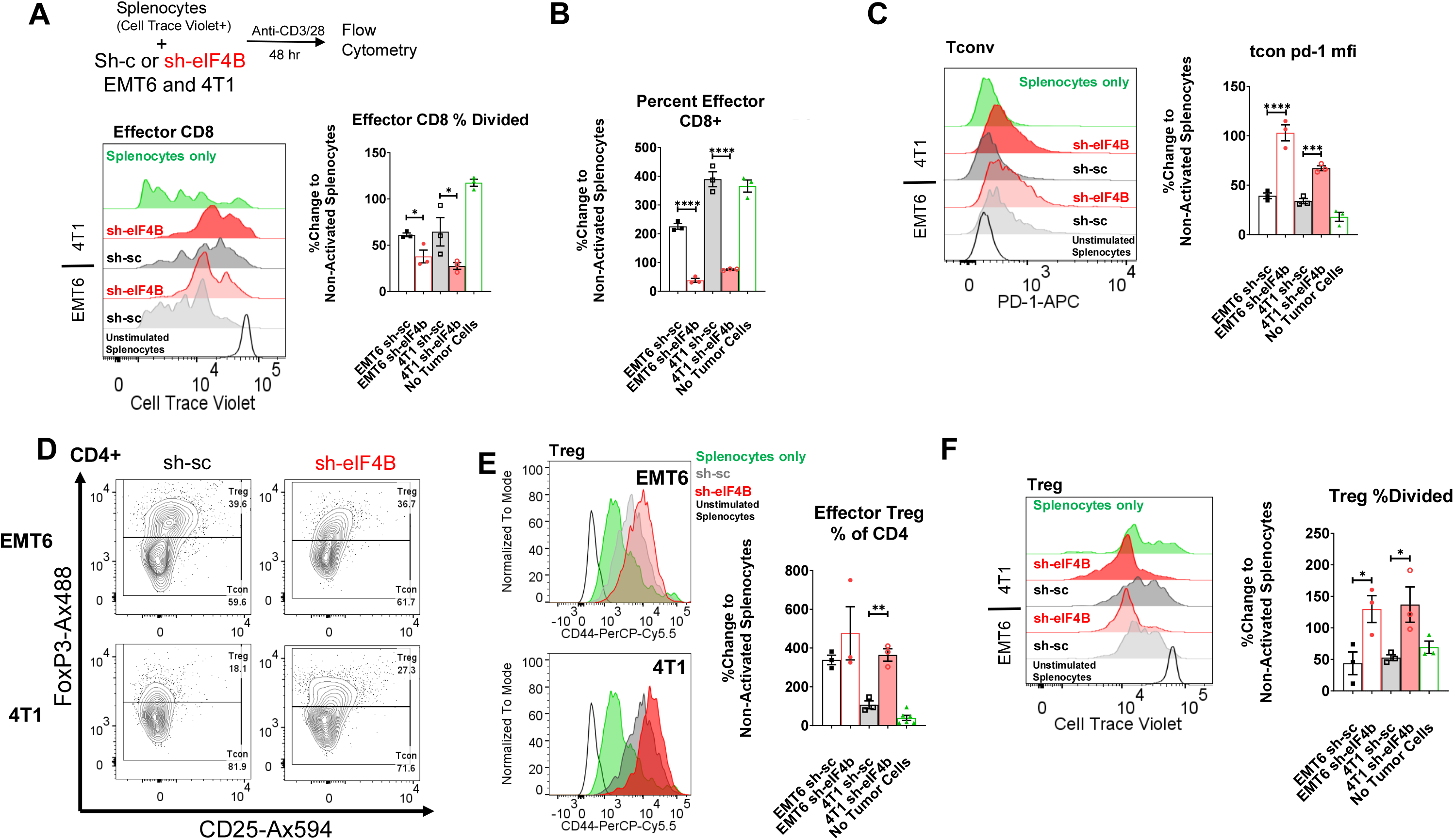
eIF4B knockdown-associated immune suppression is evident in murine tumor cell-leukocyte cocultures. Splenocyte suspensions were generated from wild type Balb/c mice, stained with proliferation-responsive fluorescent dye (Cell Trace Violet), and seeded alone in 24-well plates (1×10^6^/well), or in coculture with indicated breast cancer cell line (15,000 cells/well). Single cell type control wells and cocultures were incubated with or without CD3 and CD28 crosslinking antibodies (1ug/ml each) for 72hr prior to cell characterization by flow cytometry. Proliferation (dye dilution) by effector CD8+ T cells **(A)** as well as changes in proportions of these cells in the coculture conditions **(B)** were thus determined as were PD-1 levels on Tconv CD4+ T cells **(C)** and the frequency, activation state, and division of bulk Foxp3+ Tregs **(D)** in the cocultures. Shown are representative flow cytometry plots and corresponding quantification of key phenotypic marker changed induced by activation (mean +/-SEM). Shown are representative findings from three independent trials (\**p* < 0.05, **p<0.01, ***p<0.001, ****p<0.0001) by t test.

**Supplementary Figure S12.**
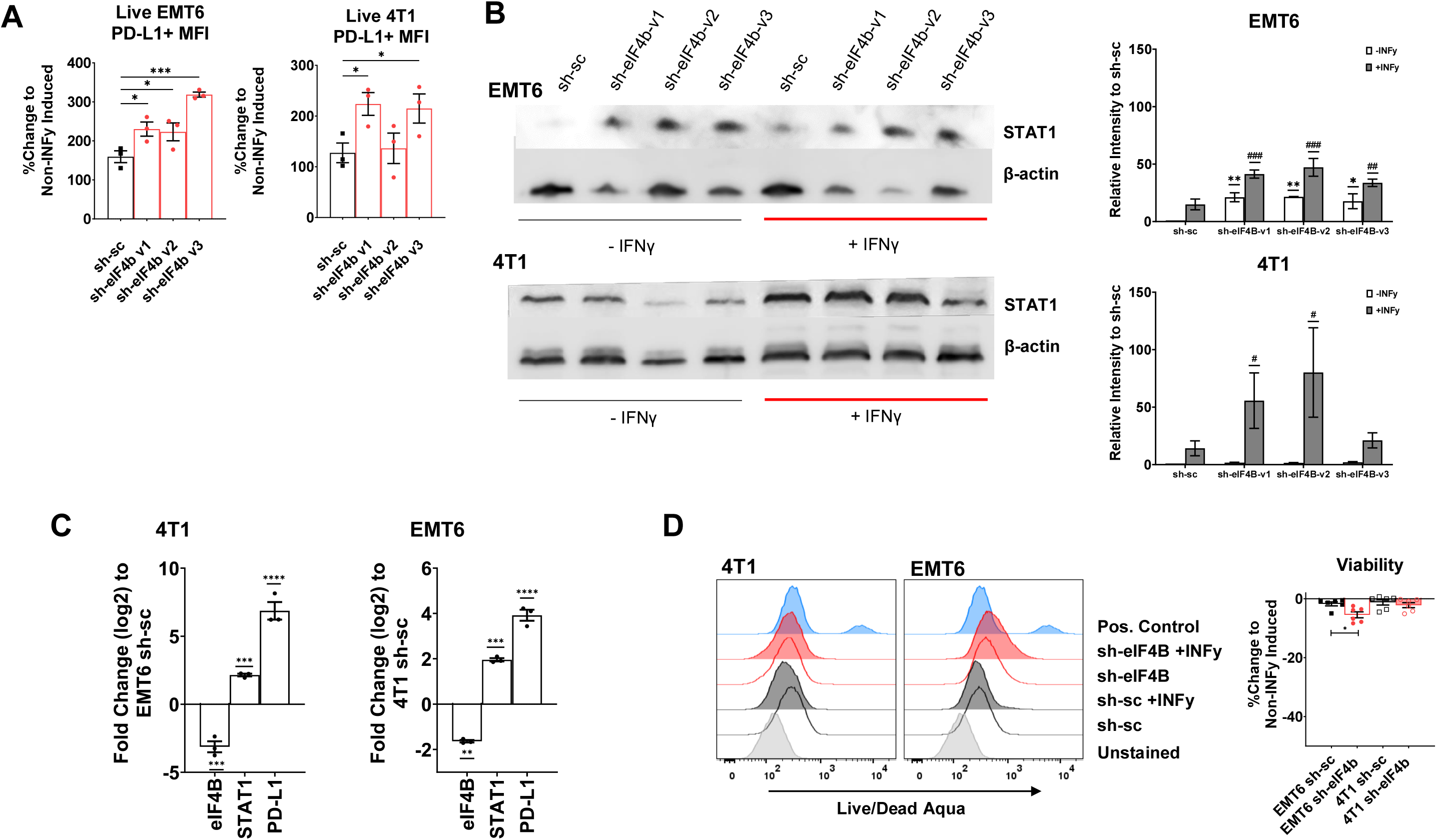
PD-L1 induction and viability in eIF4B knockdown cell lines exposed to recombinant IFN_γ_. As in the studies corresponding to Fig. 4A, murine breast cancer lines EMT6 and 4T1 stably transduced with control and sh-eIF4B constructs were grown in vitro and exposed to recombinant IFN_γ_ (100ug/ml) or media alone for 24 hours. Cells were harvested and surface PD-L1 levels were assessed by flow cytometry (**A**). Changes in PD-L1 MFI relative to media only controls were determined. Levels of STAT1 protein and transcript in these cells were found by Western Blot (beta-actin = loading control; **B**) and parallel qRT-PCR analysis, respectively, and quantified. eIF4B and PD-L1 transcript levels were also found by qRT-PCR (**C**). Tumor cell viability in these experiments was simultaneously monitored by Live/Dead Aqua staining and flow cytometry (**D**). Shown are mean values (+/-SEM) and representative images from 2-5 independent experiments. \**p* < 0.05, **p<0.01, ***p<0.001, ****p<0.0001 by t test.

**Supplementary Figure S13.**
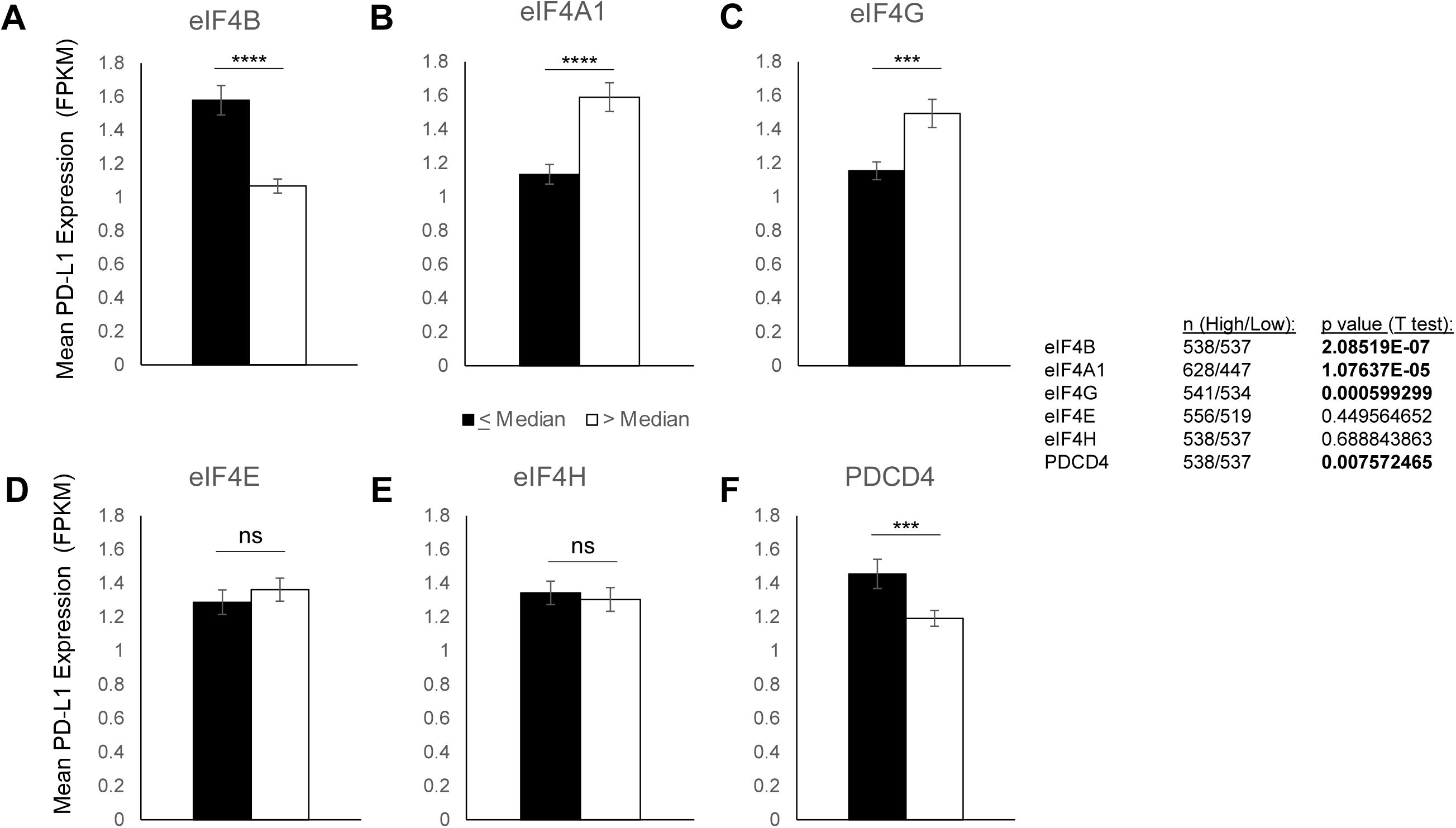
Relationships between PD-L1 expression and translation initiation factor levels in breast cancer patients. Gene expression data available for Breast Invasive Carcinoma (BRCA) patients in the TCGA (n=1075) was used to relate translation initiation factor expression to that of PD-L1. Patients were categorized by their expression of the indicated translation factor (i.e., at or below the median value or above median levels of eIF4B (A), eIF4A (B), eIF4G (C), eIF4E (D), eIF4H (E), and PDCD4 (F)). Mean levels of PD-L1 encoding transcript in each group are shown (+/-SEM). Inset table depicts patient numbers and hazard ratios for the indicated stages. \**p* < 0.05, **p<0.01, ***p<0.001, ****p<0.0001 by t test.

**Supplementary Figure S14.**
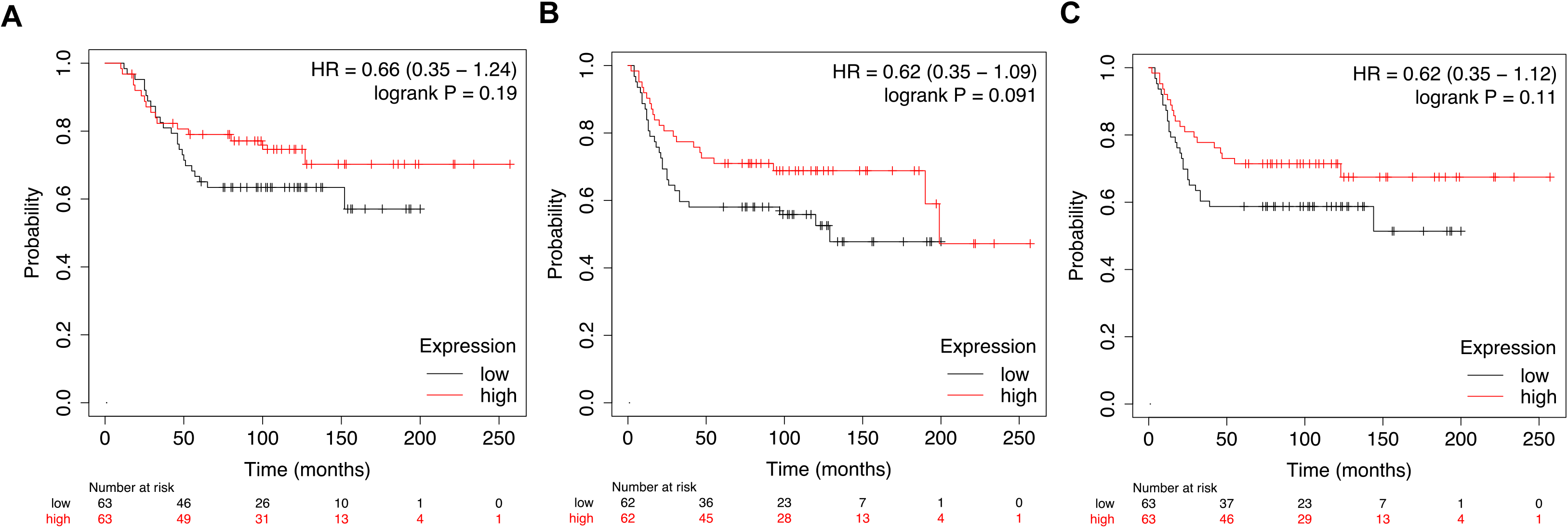
Relative eIF4B protein level and its relationship to breast cancer patient outcomes in a prior study. The association of eIF4B protein level and Overall survival (OS, **A**, n=126), recurrence free survival (RFS, **B**, n=124), and distant metastasis-free survival (DMFS, **C**, n=126), was explored using the KM plotter website and publicly available dataset previously generated by Lui et al. Shown are Kaplan-Meier survival curves and the corresponding Hazard Ratio (HR) and p values (Log Rank). Red and black lines depict high- and low-eIF4B expressing cases (above and below median expression level) respectively. An HR>1 indicates a survival disadvantage.

**Supplementary Figure S15.**
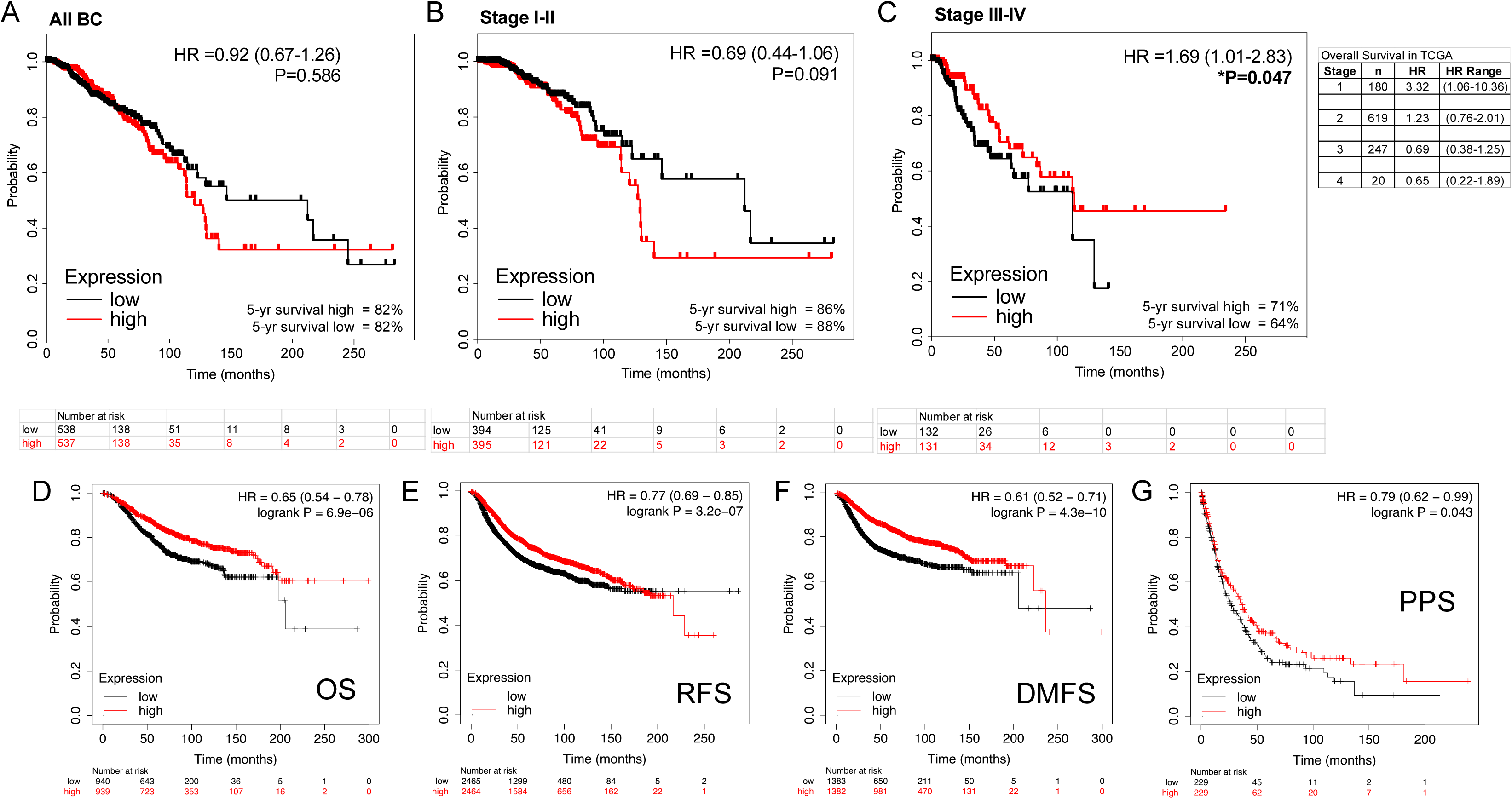
Evidence for distinct, disease stage-specific relationships between eIF4B levels and breast cancer patient outcomes. **(A-C)** Survival data for Breast Invasive Carcinoma (BRCA) patients in the TCGA was accessed through the Human Protein Atlas’ Pathology web tool. The overall survival (OS) of all patients (**A**), as well as early (stages I and II; **B**) and advanced stage patients (stage III, IV; **C**) with low (sub-median) and high (above median) eIF4B mRNA levels (FPKM) was compared. Kaplan-Meier curves were generated and the corresponding Hazard Ratio (HR) and p value for each comparison (cox regression analysis) were found. The association of *EIF4B* expression levels and the indicated survival outcomes, agnostic to any therapeutic intervention, was also explored using the KM Plotter webtool and the compiled data from at least 33 non-overlapping, publicly available gene chip (microarray) analyses. Overall survival (OS, **D**), recurrence free survival (RFS, **E**), distant metastasis-free survival (DMFS, **F**), and post-progression survival (PPS, **G**) of patients with high- and low-eIF4B expressing tumors were assessed. Red and black lines depict high- and low-*EIF4B expressing cases* (above and below median expression level) respectively. Inset tables indicates the HR associated with eIF4B high and low groups in TCGA by stage.

**Supplementary Figure S16.**
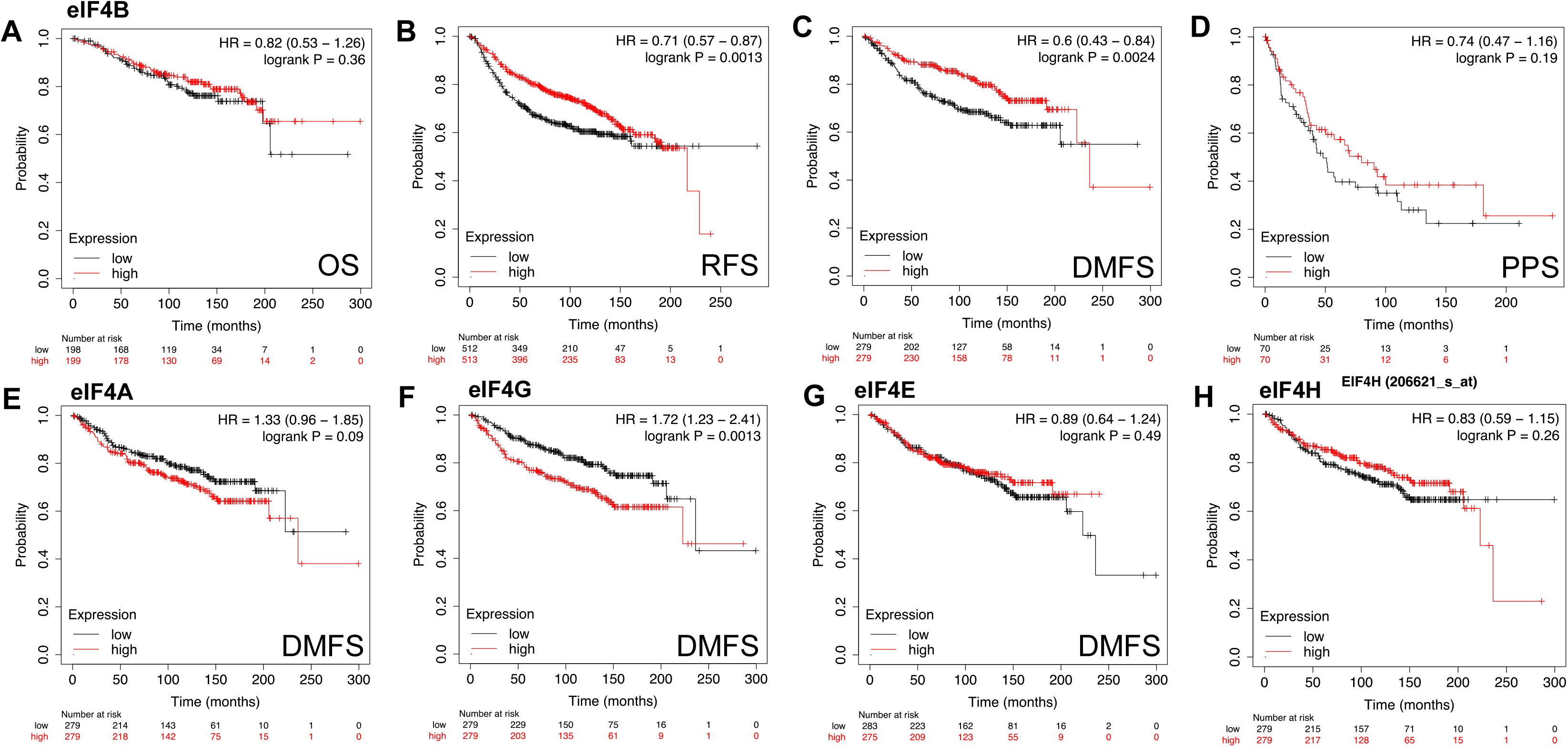
Survival outcomes for breast cancer patients expressing disparate levels of eIF4B and other translation initiation factors patients in the absence of systemic therapy. Kaplan Meier analysis of breast cancer patients stratified by eIF4B expression was performed using the KM plotter webtool and deposited, non-redundant microarray datasets. Samples from patients receiving systemic therapies were excluded from analysis. Overall survival (OS, **A**), recurrence free survival (RFS, **B**), distant metastasis-free survival (DMFS, **C**), and post-progression survival (PPS, **D**) of patients with high- and low-eIF4B expressing tumors as well as the corresponding Hazard Ratio (HR) and p value for each comparison (cox regression analysis) were determined. (**E-H**) Associations between other translation initiation factors and DMFS in these patients were similarly found. Red and black lines depict high- and low-*EIF4B expressing cases* (above and below median expression level) respectively.

**Supplementary Figure S17.**
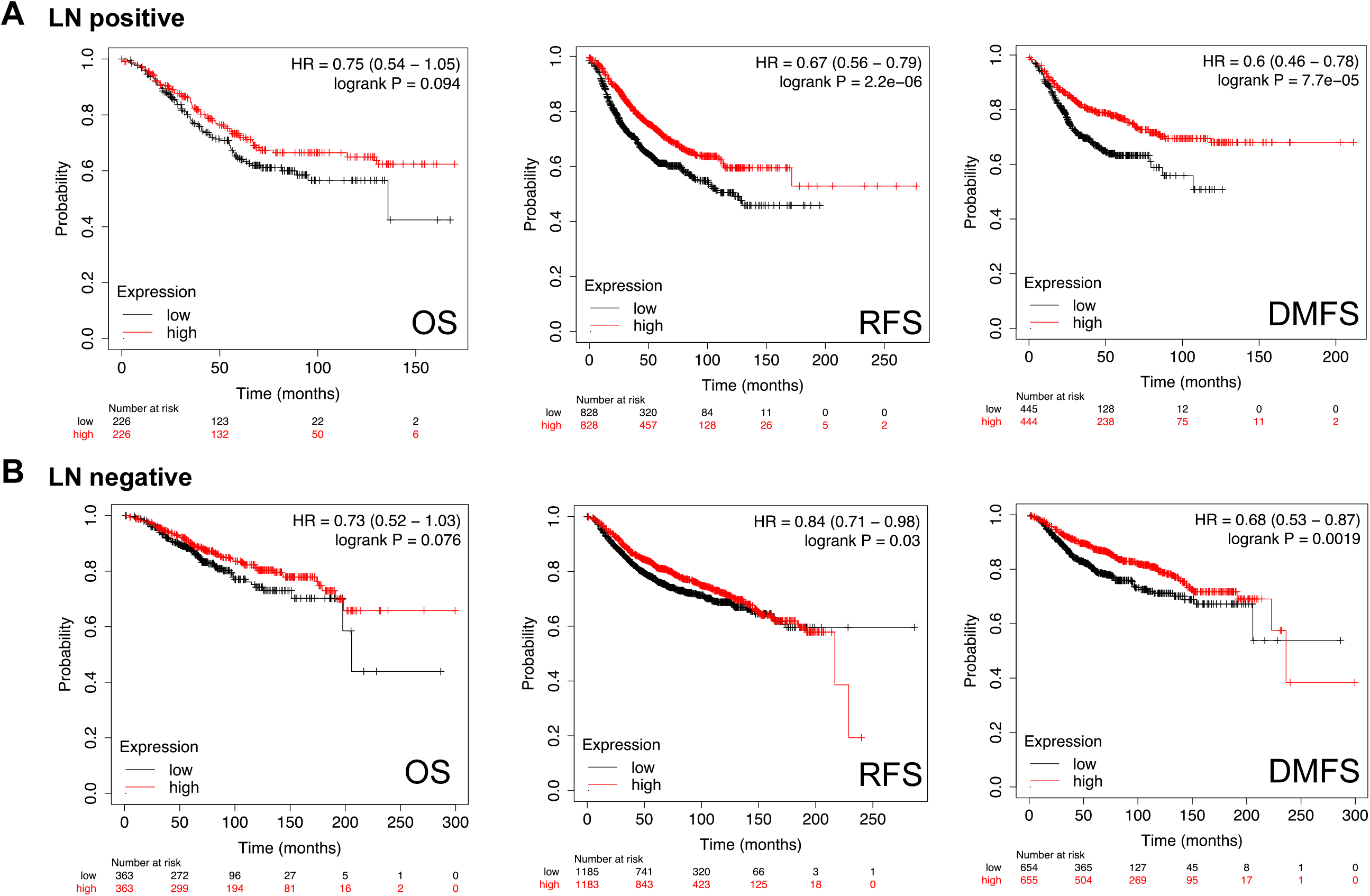
Relationship between eIF4B levels and survival outcomes in patients with and without lymph node metastasis. Kaplan Meier analysis of breast cancer patients stratified by eIF4B expression was performed using the KM plotter webtool and deposited microarray data from patients with and without lymph node metastasis. Overall survival (OS, **A**), recurrence free survival (RFS, **B**), and distant metastasis-free survival (DMFS, **C**) of patients with high- and low-eIF4B expressing tumors as well as the corresponding Hazard Ratio (HR) and p value for each comparison (cox regression analysis) were determined. Red and black lines depict high- and low-*EIF4B expressing cases* (above and below median expression level) respectively.

**Supplementary Figure S18.**
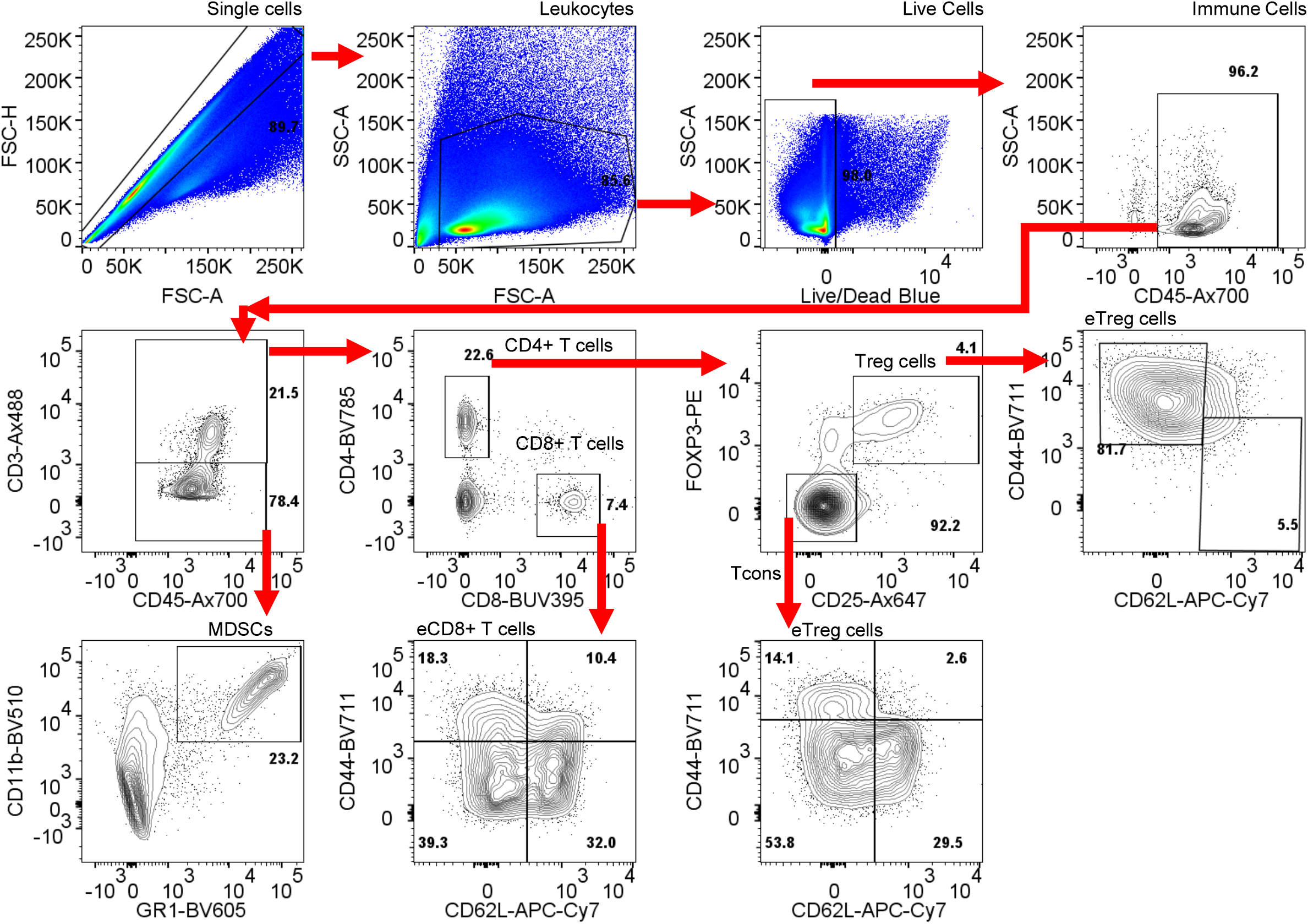
Gating strategy to identify and quantify major leukocyte subsets in flow cytometry analyses.

**Supplementary Table S1.**

| **Marker** | **Label** | **Source** | **Catalogue No.** | **Clone:** |
| --- | --- | --- | --- | --- |
| CD4 | BV785 | BioLegend | 100552 | RM4-5 |
| CD8 | BUV395 | BD Biosciences | 563786 | 53-6.7 |
| FOXP3 | AX488 | eBioscience | 53-4774-42 | 150DE4 |
| CD25 | BV605 | BioLegend | 102036 | PC61 |
| PD1 | BV421 | BioLegend | 109121 | RMP1-30 |
| LAG3 | APC | BioLegend | 125209 | C9B7W |
| TIM3 | PE | BioLegend | 134003 | B8.2C12 |
| CTLA4 | percp5.5 | BioLegend | 106316 | UC10-4B9 |
| CD45 | Ax700 | BioLegend | 103128 | 13/2.3 |
| CD44 | Ax700 | BioLegend | 103026 | IM7 |
| CD62L | APC Cy7 | BioLegend | 104428 | MEL-14 |
| Ki-67 | BV421 | BioLegend | 652411 | 16A8 |
| FOXP3 | PE | eBioscience | 12-5773-82 | FJK-16s |
| cd11b | percp5.5 | BioLegend | 101228/ | M1/70 |
| GR1 | BV605 | BioLegend | 108440 | RB6-8C5 |
| GR1 | BV605 | BD Biosciences | 563299 | RB6-8C5 |
| CD44 | BV711 | BioLegend | 103057 | IM7 |
| CD11b | BV510 | BioLegend | 101263 | M1/70 |
| Ki-67 | BV650 | BioLegend | 151215 | 11F6 |
| CD44 | PerCP-Cy5.5 | BioLegend | 103032 | IM7 |
| CD25 | Ax594 | BioLegend | 102045 | PC61 |
| PD1 | APC | BioLegend | 135209 | 29F.1A12 |
| PDL1 | PE-Cy7 | BioLegend | 124313 | 10F.9G2 |
| STAT1 | - | Cell Signaling | 9172 | - |
| eIF4B | - | Cell Signaling | 3592 | - |
| GAPDH | - | Abcam | ab8245 | - |
| beta-actin | - | Abcam | ab8226 | - |
| anti-Mouse<br>HRP | HRP | Abcam | ab205719 | - |
| anti-Rabbit<br>HRP | HRP | Abcam | ab205718 | - |

**Supplementary Table S2.**
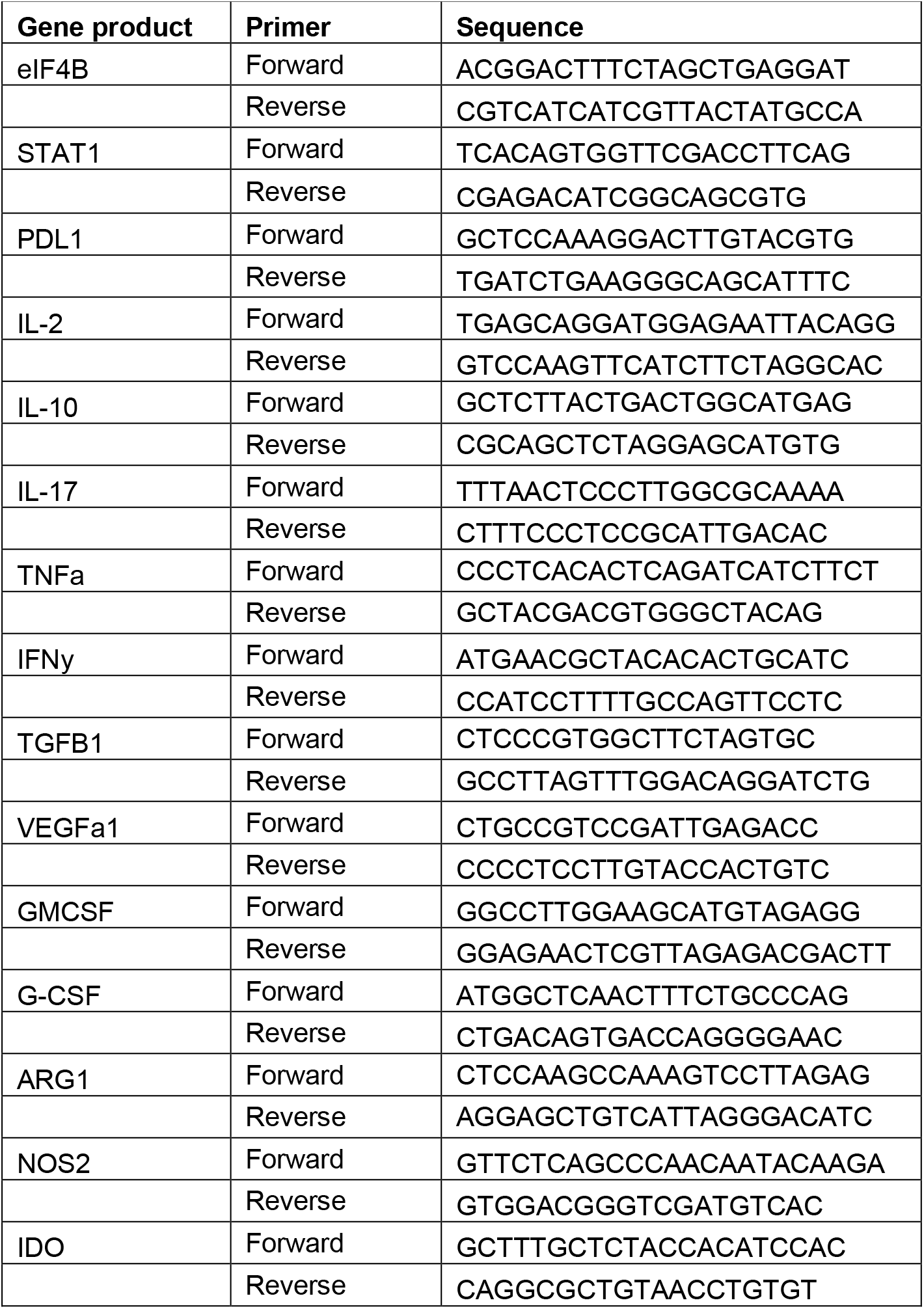

## References

1. C. Global Burden of Disease Cancer et al., Global, Regional, and National Cancer Incidence, Mortality, Years of Life Lost, Years Lived With Disability, and Disability-Adjusted Life-years for 32 Cancer Groups, 1990 to 2015: A Systematic Analysis for the Global Burden of Disease Study. JAMA Oncol 3, 524–548 (2017).

2. M. Yong et al., Survival in breast cancer patients with bone metastases and skeletal-related events: a population-based cohort study in Denmark (1999–2007). Breast Cancer Res Treat 129, 495–503 (2011).

3. C. Vaklavas, S. W. Blume, W. E. Grizzle, Translational Dysregulation in Cancer: Molecular Insights and Potential Clinical Applications in Biomarker Development. Front Oncol 7, 158 (2017).

4. J. Mladkova, M. Sanda, E. Matouskova, I. Selicharova, Phenotyping breast cancer cell lines EM-G3, HCC1937, MCF7 and MDA-MB-231 using 2-D electrophoresis and affinity chromatography for glutathione-binding proteins. BMC Cancer 10, 449 (2010).

5. J. Vydra et al., Two-dimensional electrophoretic comparison of metastatic and non-metastatic human breast tumors using in vitro cultured epithelial cells derived from the cancer tissues. BMC Cancer 8, 107 (2008).

6. M. G. Terp, R. R. Lund, O. N. Jensen, R. Leth-Larsen, H. J. Ditzel, Identification of markers associated with highly aggressive metastatic phenotypes using quantitative comparative proteomics. Cancer Genomics Proteomics 9, 265–273 (2012).

7. H. H. Milioli et al., Comparative proteomics of primary breast carcinomas and lymph node metastases outlining markers of tumor invasion. Cancer Genomics Proteomics 12, 89–101 (2015).

8. S. Wu, G. Wagner, Deep computational analysis details dysregulation of eukaryotic translation initiation complex eIF4F in human cancers. Cell Syst 12, 907–923 e906 (2021).

9. A. Modelska et al., The malignant phenotype in breast cancer is driven by eIF4A1-mediated changes in the translational landscape. Cell Death Dis 6, e1603 (2015).

10. Y. C. S. Ramon et al., Beyond molecular tumor heterogeneity: protein synthesis takes control. Oncogene 37, 2490–2501 (2018).

11. G. W. Rogers, Jr., N. J. Richter, W. F. Lima, W. C. Merrick, Modulation of the helicase activity of eIF4A by eIF4B, eIF4H, and eIF4F. J Biol Chem 276, 30914–30922 (2001).

12. L. Calviello et al., DDX3 depletion represses translation of mRNAs with complex 5' UTRs. Nucleic Acids Res 49, 5336–5350 (2021).

13. S. E. Dmitriev, I. M. Terenin, Y. E. Dunaevsky, W. C. Merrick, I. N. Shatsky, Assembly of 48S translation initiation complexes from purified components with mRNAs that have some base pairing within their 5' untranslated regions. Mol Cell Biol 23, 8925–8933 (2003).

14. T. V. Pestova, V. G. Kolupaeva, The roles of individual eukaryotic translation initiation factors in ribosomal scanning and initiation codon selection. Genes Dev 16, 2906–2922 (2002).

15. D. Maag, C. A. Fekete, Z. Gryczynski, J. R. Lorsch, A conformational change in the eukaryotic translation preinitiation complex and release of eIF1 signal recognition of the start codon. Mol Cell 17, 265–275 (2005).

16. T. Hussain et al., Structural changes enable start codon recognition by the eukaryotic translation initiation complex. Cell 159, 597–607 (2014).

17. C. A. Rubio et al., Transcriptome-wide characterization of the eIF4A signature highlights plasticity in translation regulation. Genome Biol 15, 476 (2014).

18. N. D. Sen, F. Zhou, M. S. Harris, N. T. Ingolia, A. G. Hinnebusch, eIF4B stimulates translation of long mRNAs with structured 5' UTRs and low closed-loop potential but weak dependence on eIF4G. Proc Natl Acad Sci U S A 113, 10464–10472 (2016).

19. E. H. Park, F. Zhang, J. Warringer, P. Sunnerhagen, A. G. Hinnebusch, Depletion of eIF4G from yeast cells narrows the range of translational efficiencies genome-wide. BMC Genomics 12, 68 (2011).

20. S. Iwasaki et al., The Translation Inhibitor Rocaglamide Targets a Bimolecular Cavity between eIF4A and Polypurine RNA. Mol Cell 73, 738–748 e739 (2019).

21. H. S. Yang et al., The transformation suppressor Pdcd4 is a novel eukaryotic translation initiation factor 4A binding protein that inhibits translation. Mol Cell Biol 23, 26–37 (2003).

22. M. Oberer, A. Marintchev, G. Wagner, Structural basis for the enhancement of eIF4A helicase activity by eIF4G. Genes Dev 19, 2212–2223 (2005).

23. N. Robichaud et al., Translational control in the tumor microenvironment promotes lung metastasis: Phosphorylation of eIF4E in neutrophils. Proc Natl Acad Sci U S A 115, E2202–E2209 (2018).

24. N. Robichaud, N. Sonenberg, D. Ruggero, R. J. Schneider, Translational Control in Cancer. Cold Spring Harb Perspect Biol 10.1101/cshperspect.a032896 (2018).

25. Y. Chen et al., Loss of PDCD4 expression in human lung cancer correlates with tumour progression and prognosis. J Pathol 200, 640–646 (2003).

26. E. Kuang, B. Fu, Q. Liang, J. Myoung, F. Zhu, Phosphorylation of eukaryotic translation initiation factor 4B (EIF4B) by open reading frame 45/p90 ribosomal S6 kinase (ORF45/RSK) signaling axis facilitates protein translation during Kaposi sarcoma-associated herpesvirus (KSHV) lytic replication. J Biol Chem 286, 41171–41182 (2011).

27. D. Shahbazian et al., The mTOR/PI3K and MAPK pathways converge on eIF4B to control its phosphorylation and activity. EMBO J 25, 2781–2791 (2006).

28. B. Raught et al., Phosphorylation of eucaryotic translation initiation factor 4B Ser422 is modulated by S6 kinases. EMBO J 23, 1761–1769 (2004).

29. A. De Benedetti, J. R. Graff, eIF-4E expression and its role in malignancies and metastases. Oncogene 23, 3189–3199 (2004).

30. V. Kerekatte et al., The proto-oncogene/translation factor eIF4E: a survey of its expression in breast carcinomas. Int J Cancer 64, 27–31 (1995).

31. X. Liu, H. Moshiri, S. E. Walker, Binding to the Ribosome by Eukaryotic Initiation Factor 4B Drives Yeast Translational Control in Response to Urea. bioRXIV http://dx.doi.org/10.1101/819672 (2019).

32. T. M. Goldson et al., Eukaryotic initiation factor 4E variants alter the morphology, proliferation, and colony-formation properties of MDA-MB-435 cancer cells. Mol Carcinog 46, 71–84 (2007).

33. B. D. Li, L. Liu, M. Dawson, A. De Benedetti, Overexpression of eukaryotic initiation factor 4E (eIF4E) in breast carcinoma. Cancer 79, 2385–2390 (1997).

34. N. Methot, A. Pause, J. W. Hershey, N. Sonenberg, The translation initiation factor eIF-4B contains an RNA-binding region that is distinct and independent from its ribonucleoprotein consensus sequence. Mol Cell Biol 14, 2307–2316 (1994).

35. S. E. Walker et al., Yeast eIF4B binds to the head of the 40S ribosomal subunit and promotes mRNA recruitment through its N-terminal and internal repeat domains. RNA 19, 191–207 (2013).

36. A. R. Ozes, K. Feoktistova, B. C. Avanzino, C. S. Fraser, Duplex unwinding and ATPase activities of the DEAD-box helicase eIF4A are coupled by eIF4G and eIF4B. J Mol Biol 412, 674–687 (2011).

37. A. L. Wolfe et al., RNA G-quadruplexes cause eIF4A-dependent oncogene translation in cancer. Nature 513, 65–70 (2014).

38. C. M. Howard et al., The CXCR4-LASP1-eIF4F Axis Promotes Translation of Oncogenic Proteins in Triple-Negative Breast Cancer Cells. Front Oncol 9, 284 (2019).

39. O. Attar-Schneider, L. Drucker, M. Gottfried, Migration and epithelial-to-mesenchymal transition of lung cancer can be targeted via translation initiation factors eIF4E and eIF4GI. Lab Invest 96, 1004–1015 (2016).

40. D. Shahbazian et al., Control of cell survival and proliferation by mammalian eukaryotic initiation factor 4B. Mol Cell Biol 30, 1478–1485 (2010).

41. E. Horvilleur et al., A role for eukaryotic initiation factor 4B overexpression in the pathogenesis of diffuse large B-cell lymphoma. Leukemia 28, 1092–1102 (2014).

42. C. Gong et al., Phosphorylation independent eIF4E translational reprogramming of selective mRNAs determines tamoxifen resistance in breast cancer. Oncogene 10.1038/s41388-020-1210-y (2020).

43. S. J. Park, B. H. Yoon, S. K. Kim, S. Y. Kim, GENT2: an updated gene expression database for normal and tumor tissues. BMC Med Genomics 12, 101 (2019).

44. M. Uhlen et al., A pathology atlas of the human cancer transcriptome. Science 357 (2017).

45. Z. Zhang et al., Global analysis of tRNA and translation factor expression reveals a dynamic landscape of translational regulation in human cancers. Commun Biol 1, 234 (2018).

46. F. M. Howard, O. I. Olopade, Epidemiology of Triple-Negative Breast Cancer: A Review. Cancer J 27, 8–16 (2021).

47. M. Ouzounova et al., Monocytic and granulocytic myeloid derived suppressor cells differentially regulate spatiotemporal tumour plasticity during metastatic cascade. Nat Commun 8, 14979 (2017).

48. R. Piranlioglu et al., Primary tumor-induced immunity eradicates disseminated tumor cells in syngeneic mouse model. Nat Commun 10, 1430 (2019).

49. M. Ghandi et al., Next-generation characterization of the Cancer Cell Line Encyclopedia. Nature 569, 503–508 (2019).

50. B. I. Bassey-Archibong et al., Kaiso depletion attenuates transforming growth factor-beta signaling and metastatic activity of triple-negative breast cancer cells. Oncogenesis 5, e208 (2016).

51. E. Iorns et al., A new mouse model for the study of human breast cancer metastasis. PLoS One 7, e47995 (2012).

52. A. Herrera-Gayol, S. Jothy, CD44 modulates Hs578T human breast cancer cell adhesion, migration, and invasiveness. Exp Mol Pathol 66, 99–108 (1999).

53. N. Tasdemir et al., Comprehensive Phenotypic Characterization of Human Invasive Lobular Carcinoma Cell Lines in 2D and 3D Cultures. Cancer Res 78, 6209–6222 (2018).

54. T. H. Lee et al., Vascular endothelial growth factor mediates intracrine survival in human breast carcinoma cells through internally expressed VEGFR1/FLT1. PLoS Med 4, e186 (2007).

55. Y. C. Liao et al., Overexpressed hPTTG1 promotes breast cancer cell invasion and metastasis by regulating GEF-H1/RhoA signalling. Oncogene 31, 3086–3097 (2012).

56. Y. Wang et al., Mitotic MELK-eIF4B signaling controls protein synthesis and tumor cell survival. Proc Natl Acad Sci U S A 113, 9810–9815 (2016).

57. E. K. Schmidt, G. Clavarino, M. Ceppi, P. Pierre, SUnSET, a nonradioactive method to monitor protein synthesis. Nat Methods 6, 275–277 (2009).

58. L. E. Barney et al., A cell-ECM screening method to predict breast cancer metastasis. Integr Biol (Camb) 7, 198–212 (2015).

59. C. V. Hinton, S. Avraham, H. K. Avraham, Contributions of integrin-linked kinase to breast cancer metastasis and tumourigenesis. J Cell Mol Med 12, 1517–1526 (2008).

60. J. Schaller, J. Agudo, Metastatic Colonization: Escaping Immune Surveillance. Cancers (Basel) 12 (2020).

61. M. Mohme, S. Riethdorf, K. Pantel, Circulating and disseminated tumour cells-mechanisms of immune surveillance and escape. Nat Rev Clin Oncol 14, 155–167 (2017).

62. J. L. Chao, P. A. Savage, Unlocking the Complexities of Tumor-Associated Regulatory T Cells. J Immunol 200, 415–421 (2018).

63. F. Mazerolles, F. Rieux-Laucat, PD-L1 is expressed on human activated naive effector CD4+ T cells. Regulation by dendritic cells and regulatory CD4+ T cells. PLoS One 16, e0260206 (2021).

64. M. De Simone et al., Transcriptional Landscape of Human Tissue Lymphocytes Unveils Uniqueness of Tumor-Infiltrating T Regulatory Cells. Immunity 45, 1135–1147 (2016).

65. D. I. Gabrilovich, S. Nagaraj, Myeloid-derived suppressor cells as regulators of the immune system. Nat Rev Immunol 9, 162–174 (2009).

66. M. Cerezo et al., Translational control of tumor immune escape via the eIF4F-STAT1-PD-L1 axis in melanoma. Nat Med 24, 1877–1886 (2018).

67. L. M. Francisco et al., PD-L1 regulates the development, maintenance, and function of induced regulatory T cells. J Exp Med 206, 3015–3029 (2009).

68. N. Q. Liu et al., Comparative proteome analysis revealing an 11-protein signature for aggressive triple-negative breast cancer. J Natl Cancer Inst 106, djt376 (2014).

69. K. Chin et al., Genomic and transcriptional aberrations linked to breast cancer pathophysiologies. Cancer Cell 10, 529–541 (2006).

70. P. Jezequel et al., Gene-expression molecular subtyping of triple-negative breast cancer tumours: importance of immune response. Breast Cancer Res 17, 43 (2015).

71. K. Chan et al., eIF4A supports an oncogenic translation program in pancreatic ductal adenocarcinoma. Nat Commun 10, 5151 (2019).

72. C. Xue, X. Gu, G. Li, Z. Bao, L. Li, Expression and Functional Roles of Eukaryotic Initiation Factor 4A Family Proteins in Human Cancers. Front Cell Dev Biol 9, 711965 (2021).

73. D. Shahbazian, A. Parsyan, E. Petroulakis, J. Hershey, N. Sonenberg, eIF4B controls survival and proliferation and is regulated by proto-oncogenic signaling pathways. Cell Cycle 9, 4106–4109 (2010).

74. S. Fan et al., Phosphorylated eukaryotic translation initiation factor 4 (eIF4E) is elevated in human cancer tissues. Cancer Biol Ther 8, 1463–1469 (2009).

75. B. Hellwig et al., Epsin Family Member 3 and Ribosome-Related Genes Are Associated with Late Metastasis in Estrogen Receptor-Positive Breast Cancer and Long-Term Survival in Non-Small Cell Lung Cancer Using a Genome-Wide Identification and Validation Strategy. PLoS One 11, e0167585 (2016).

76. S. Wu, G. Wagner, Deep computational analysis of human cancer and non-cancer tissues details dysregulation of eIF4F components and their interactions in human cancers. bioRXiV https://doi.org/10.1101/2020.10.12.336263 (2020).

77. M. S. Holtz, S. J. Forman, R. Bhatia, Nonproliferating CML CD34+ progenitors are resistant to apoptosis induced by a wide range of proapoptotic stimuli. Leukemia 19, 1034–1041 (2005).

78. T. Naranda, W. B. Strong, J. Menaya, B. J. Fabbri, J. W. Hershey, Two structural domains of initiation factor eIF-4B are involved in binding to RNA. J Biol Chem 269, 14465–14472 (1994).

79. N. Rozovsky, A. C. Butterworth, M. J. Moore, Interactions between eIF4AI and its accessory factors eIF4B and eIF4H. RNA 14, 2136–2148 (2008).

80. S. Suresh, K. A. O'Donnell, Translational Control of Immune Evasion in Cancer. Trends Cancer 7, 580–582 (2021).

81. S. Suresh et al., eIF5B drives integrated stress response-dependent translation of PD-L1 in lung cancer. Nat Cancer 1, 533–545 (2020).

82. F. Li et al., High expression of eIF4E is associated with tumor macrophage infiltration and leads to poor prognosis in breast cancer. BMC Cancer 21, 1305 (2021).

83. M. Mazel et al., Frequent expression of PD-L1 on circulating breast cancer cells. Mol Oncol 9, 1773–1782 (2015).

84. S. Bergmann et al., Evaluation of PD-L1 expression on circulating tumor cells (CTCs) in patients with advanced urothelial carcinoma (UC). Oncoimmunology 9, 1738798 (2020).

85. D. Kong et al., Correlation between PD-L1 expression ON CTCs and prognosis of patients with cancer: a systematic review and meta-analysis. Oncoimmunology 10, 1938476 (2021).

86. Y. Wang et al., PD-L1 Expression in Circulating Tumor Cells Increases during Radio(chemo)therapy and Indicates Poor Prognosis in Non-small Cell Lung Cancer. Sci Rep 9, 566 (2019).

87. M. Rozenblit et al., Comparison of PD-L1 protein expression between primary tumors and metastatic lesions in triple negative breast cancers. J Immunother Cancer 8 (2020).

88. C. Boman et al., Discordance of PD-L1 status between primary and metastatic breast cancer: A systematic review and meta-analysis. Cancer Treat Rev 99, 102257 (2021).

89. C. Yuan et al., Expression of PD-1/PD-L1 in primary breast tumours and metastatic axillary lymph nodes and its correlation with clinicopathological parameters. Sci Rep 9, 14356 (2019).

90. M. Li et al., Heterogeneity of PD-L1 expression in primary tumors and paired lymph node metastases of triple negative breast cancer. BMC Cancer 18, 4 (2018).

91. A. Albini, R. Benelli, The chemoinvasion assay: a method to assess tumor and endothelial cell invasion and its modulation. Nat Protoc 2, 504–511 (2007).

92. A. Nagy, G. Munkacsy, B. Gyorffy, Pancancer survival analysis of cancer hallmark genes. Sci Rep 11, 6047 (2021).

93. A. Osz, A. Lanczky, B. Gyorffy, Survival analysis in breast cancer using proteomic data from four independent datasets. Sci Rep 11, 16787 (2021).

94. B. Gyorffy, Survival analysis across the entire transcriptome identifies biomarkers with the highest prognostic power in breast cancer. Comput Struct Biotechnol J 19, 4101–4109 (2021).

95. Q. Li, N. J. Birkbak, B. Gyorffy, Z. Szallasi, A. C. Eklund, Jetset: selecting the optimal microarray probe set to represent a gene. BMC Bioinformatics 12, 474 (2011).

